# Maternal style does not predict infant growth and survival in wild Guinea baboons

**DOI:** 10.64898/2026.01.09.698613

**Authors:** Anaïs Avilés de Diego, Federica Dal Pesco, Roger Mundry, Julia Fischer

**Affiliations:** Department of Primate Cognition, Georg-August-Universität Göttingen, Johann-Friedrich Blumenbach Institute, Göttingen, Germany; Cognitive Ethology Laboratory, German Primate Center, Göttingen, Germany; Leibniz ScienceCampus Primate Cognition, Göttingen, Germany

**Keywords:** Growth, mother-infant relationship, maternal style, *Papio papio*, Parallel Laser Photogrammetry, survival

## Abstract

Primate infants depend on their mothers for nursing, transportation, and survival, but also benefit from other behaviours such as contact, protection, and the mother serving as a model for social learning. During infancy, variation in maternal care (“maternal style”) can have profound consequences for infant development and survival. We investigated the link between maternal style and infant growth and survival in wild Guinea baboons (*Papio papio*). This species lives in a tolerant social system with female-biased dispersal. We used data from *N* = 80 infants to assess variation in maternal style. The final statistical models comprised data from *N* = 50 infants for the growth analysis based on Parallel Laser Photogrammetry (PLP) measures of forearm length, and *N* = 63 infants for the survival analysis. To account for heterogeneity in data, we applied a recently established statistical framework that controls for variation in maternal behaviour with infant age. We found only moderate variation among mothers and no evidence for a link between maternal style and infant growth or survival. Neither infant sex, maternal age, nor NDVI variation, a measure later shown to be a poor indicator of food availability, predicted infant growth or survival. We suspect that, in this population living in a habitat with high carrying capacity, disease and predation have greater effects on infant growth and survival than maternal behaviour. Future studies should consider the interplay between infant behaviour and maternal responses in greater detail to better understand variation in early social development.

## Introduction

In many mammalian species, mothers have a profound influence on their infants’ growth and survival, providing essential nutrition, social support, and protection (e.g., subantarctic fur seal, *Arctocephalus tropicalis*: Georges & Guinet, 2000; bottlenose dolphins, *Tursiops truncatus*: Hill et al., 2007; Mann & Watson-Capps, 2005; African elefants, *Loxodonta africana*: Lee & Moss, 1986). In primates, the mother-infant bond is particularly important, and mothers not only nurse and protect their offspring but also carry them. Mothers also serve as models for social learning (Whiten & van de Waal, 2018) and constitute crucial anchors during social development (van Boekholt et al., 2021). Individual differences in how mothers behave towards their infants are known as maternal style (Altmann, 1980; Fairbanks, 1996; Nicolson, 1987) and have been related to the mother’s personality, age, and parity (reviewed in Fairbanks, 1996). Studies of maternal style typically describe maternal behaviours along two dimensions: protectiveness and rejection (e.g., yellow baboons, *Papio cynocephalus*: Altmann, 1980; rhesus macaques, *Macaca mulatta*: Berman, 1990a, 1990b; vervet monkeys, *Chlorocebus pygerythrus*: Fairbanks & McGuire, 1987). Protective mothers invest more in caregiving, in maintaining contact with their infants, and in restricting their infants’ explorations. Rejecting mothers are more tolerant of separation, less attentive, allow more interactions with other group members than protective mothers, and are more likely to increase the distance to the infant.

Infants of rejecting mothers experience less contact earlier (Altmann, 1980), exhibit more exploratory behaviour, and are less afraid of novelty (Bardi & Huffman, 2002; Fairbanks & McGuire, 1988). Yet, a rejecting maternal style might also be associated with higher infant mortality. In yellow baboons, infants of rejecting mothers experienced slightly higher rates of illness and death (Altmann, 1980): five out of seven infants born to rejecting (here termed “laissez-faire”) mothers became ill or died, while two out of five infants born to protective (here termed “restrictive”) mothers became ill or died. In vervet monkeys, infant mortality was more pronounced among infants with lower suckling success and early weaning (Lee, 1984), whereas a more protective maternal style was associated with higher survival in captive vervet monkeys (Fairbanks, 1996).

Maternal behaviour is supposed to vary with age and experience. The parental investment theory (Trivers, 1972, 1974) predicts that younger mothers should terminate investment in their offspring earlier because they have more to gain from future reproduction than older mothers. Indeed, in wild mandrills, *Mandrillus sphinx*, older mothers spent more time in contact with their infants and groomed them more than younger mothers (Roura-Torres et al., 2025), and in captive gorillas, *Gorilla gorilla gorilla*, more experienced mothers spent more time in contact with younger than with older offspring (Amici et al., 2025). In Barbary macaques, *Macaca sylvanus*, older mothers spent more time in contact with their offspring than young mothers, supporting the parental investment theory (Paul et al., 1993). The maternal experience hypothesis, by contrast, posits that young mothers should be more protective than older mothers because they are less experienced. Evidence in favour of the maternal experience hypothesis came from a study of rhesus macaques (Holley & Simpson, 1981), in which primiparous mothers were less confident than multiparous mothers, and from a study with Japanese macaques, *Macaca fuscata*, (Schino et al., 1995), in which multiparous mothers were less protective than primiparous ones. Evidence from other mammal species also highlights the role of experience in shaping maternal behaviour. In horses (*Equus caballus*, Cameron et al., 2000), older mares were more protective during the first weeks after birth, when foal survival is most critical, but reduced maternal diligence later and were more likely to reproduce in subsequent years, suggesting that experienced mothers may allocate care more effectively rather than increasing overall maternal investment.

Furthermore, social, demographic, and ecological variables may be linked to maternal style. For example, a low maternal dominance rank (Altmann, 1980), the presence of newly immigrated males (Fairbanks & McGuire, 1987), or group members who “steal” the infant (Rowell et al., 1964) may increase maternal protectiveness. Conversely, mothers may be more rejecting when surrounded by close kin that offer social support (Altmann, 1980; Berman, 1980). Additionally, the quality and availability of food, which in turn impact maternal health, are crucial. When food is limited, mothers might be unable to sustain milk production at levels that benefit offspring survival, as shown in African elephants, in which low food availability during drought years constrains lactation and affects calf survival (Lee & Moss, 1986). In such conditions, lactating mothers should wean their infants earlier and therefore be more rejecting (Clutton-Brock, 1991). External risk variables may also increase protectiveness; for example, rhesus macaque mothers became more protective after trapping activities on Cayo Santiago (Berman, 1989), and bottlenose dolphin mothers altered their behaviour in response to predation risk (Mann & Watson-Capps, 2005).

In recent years, differences in maternal style and the consequences for offspring have attracted increasing attention. Revathe and colleagues (2025) investigated individual differences among Sumatran orangutan mothers, *Pongo abelii*, using longitudinal and cross-sectional behavioural data from 15 mothers in the Suaq Balimbing research area. Mothers differed substantially and consistently in four of the six maternal behaviours studied: contact termination, close-proximity termination, carrying, and feeding in proximity. Rolland and colleagues (2025) studied offspring responses to natural threats to infer variation in attachment in wild western chimpanzees, *Pan troglodytes verus*. They observed considerable differences among offspring and used these to assign attachment types. Fedurek and colleagues (2025) compared mother– infant proximity in two wild chimpanzee populations (eastern chimpanzees, *Pan troglodytes schweinfurthii*, and western chimpanzees) with varying levels of infanticide risk. In the study population with the higher infanticide risk, less-gregarious mothers stayed closer to their infants when many other adult females were present; the opposite pattern held for highly gregarious mothers. These studies point to the intricate interplay between maternal personality and infant experiences in primate species. Yet, how maternal style affects growth and survival is less well understood, and previous analyses lacked the statistical means to control for variation in maternal investment with respect to infant age (Mundry et al., 2023).

Here, we investigate how differences in maternal style influence the growth and survival of wild infant Guinea baboons (*Papio papio*). Guinea baboons provide insight into how the social environment influences the mother-infant relationship, as they live in a complex multi-level society (Fischer et al., 2017; Patzelt et al., 2014) characterized by low levels of both within- and between-group aggression. Genetic and behavioural evidence indicate that Guinea baboons exhibit female-biased dispersal (Kopp et al., 2015), with females transferring relatively freely between different societal levels (Goffe et al., 2016). To test the link between maternal style and infant growth and survival, we first characterized maternal style. We predicted that infants who spend less time and have fewer interactions with their mothers have slower growth trajectories and experience higher mortality than those who associate and interact more with their mothers. We included maternal age as a proxy for cumulative maternal experience, given evidence that maternal experience, particularly early in the reproductive lifespan, can affect an infant’s growth (e.g., Altmann & Alberts, 2005) and survival (e.g., Arlet et al., 2014; Pusey, 2012). Further, we included infant sex to control for potential sex differences in growth and survival (e.g., due to early emergence of sexual dimorphism). We also controlled for unit size following the conjecture that females may compete over access to the primary male for infant protection. Finally, we controlled for variation in ecological conditions.

One challenge when analysing the link between maternal style and infant growth and survival is controlling for variation in maternal behaviour with infant age. In brief, at any given point in time, the variation in maternal behaviour could be due to variation between mothers or due to variation in infant age. Failure to control for infant age can give rise to spurious results (Mundry et al., 2023). Moreover, longitudinal data, particularly from wild animal populations, are often highly heterogeneous. Animals disappear or die, and it may be impossible to continuously monitor study groups. We therefore employed an analytical framework that enabled us to account for changes in behaviour with infant age and data gaps when analysing ontogenetic processes (Mundry et al., 2023). We also assessed the repeatability of maternal behaviour. Finally, we conducted an exploratory analysis to examine whether maternal age or infant sex directly affected maternal style.

## Methods

### Compliance with ethical standards

We followed all applicable international, national, and/or institutional guidelines for the care and use of animals. Approval and research permission were granted by the Direction des Parcs Nationaux (DPN) and the Ministère de l’Environnement et de la Protection de la Nature (MEPN) de la République du Sénégal. This research complies with the regulations set by the Senegalese agencies and the Animal Care Committee at the German Primate Centre (Germany). This study followed the ASAB guidelines for the ethical treatment of nonhuman animals in behavioural research (ASAB Ethical Committee/ABS Animal Care Committee, 2023).

#### Study subjects and field site

We collected behavioural data from Guinea baboons ranging near the Centre de Recherche de Primatologie (CRP) Simenti in the Niokolo-Koba National Park, Senegal (described in Fischer et al., 2017). Guinea baboons live in a multi-level social organisation, with ‘units’ comprising one reproductively active ‘primary’ male, one to six females, and their immature offspring (Dal Pesco et al., 2022; Goffe et al., 2016). Primary males largely monopolize matings and conceptions within their units (Dal Pesco et al., 2022; Goffe et al., 2016). Several units associate to form a ‘party’, and two or more parties group together into ‘gangs’ (Patzelt et al., 2014). Gangs have overlapping home ranges, and competition over food or sleeping trees is low (Ohrndorf et al., 2025a, 2025b; Patzelt et al., 2014). Senegal has a pronounced seasonality: a rainy season from June to October and a dry season from November to April, with an average annual precipitation of 956 mm (Zinner et al., 2021). Seasonal variation in rainfall is related to plant productivity. The study population’s home range and daily travel distances increase in the rainy season (Zinner et al., 2021).

We collected behavioural data from April 2017 to December 2021 and growth data using Parallel Laser Photogrammetry (PLP) from April 2019 to August 2021. We used demographic records from long-term data to assess birth dates, maternal age, unit size, and survival. The study subjects were all infants up to 18 months of age associated with the study parties during the study period. We began data collection on two parties, party 5 and party 6, in April 2017 and expanded to additional parties over time, adding party 9 in December 2018 and parties 13 and 15 in February 2021. During the study period, several parties underwent fissions (e.g., party 9 split to form 9B in 2019, and party 6 split into 6W and 6I in 2020). We continued monitoring the resulting parties thereafter. We conducted observations and recorded demographic information daily from 06:30 to 13:00 hours. We identified all group members by their physical characteristics. To record the data, we used handheld devices (Samsung Galaxy Note II GT-7100 or Gigaset GX290) and the Pendragon software version 7.2.21 (Pendragon Software Corporation, Chicago, IL, USA). Observer reliability and identification accuracy were assessed before observations.

Our dataset is characterized by many subjects but significant heterogeneity (Fig. S1). Overall, 11 infants were born before the study started, while 20 infants were still younger than 18-months when the study ended. In addition, researchers left the field site due to the COVID-19 pandemic from April to November 2020. Six infants were born during the researchers’ absence. These individuals were subsequently included in the study and assigned ages based on developmental changes in natal coat colour (from black to brown) and skin pigmentation (from light to dark). The pandemic interruption led to heterogeneous sampling histories among individuals. Eleven infants experienced a gap in data collection during their first 18 months of life, while eight were not resampled after the interruption due to their transition into older age classes or absence from the study party upon return. Overall, we sampled 80 infants (45 females) born to 49 mothers. Mothers had at least one and up to a maximum of four infants during the study period (mean = 1.63, SD= 0.92), with 39% of mothers giving birth to multiple infants.

Seventeen infants had young adult mothers, 50 had middle-aged mothers, and 13 had old mothers (Dal Pesco & Fischer, 2022); 10 mothers with more than one infant were assigned to different age categories. Exact ages were unknown for most mothers (94%) because systematic individual monitoring had only started in 2011. In addition, in this species, dispersal is female-biased, leading to the immigration of females with unknown birth dates into our study parties. We therefore assigned females’ age using standardized morphological and developmental categories that incorporate both physical characteristics and reproductive milestones (see Dal Pesco & Fischer, 2022). As a result, maternal age category (young, middle-aged, old) and parity status (primiparous, multiparous) were highly correlated. All primiparous females were classified as young, whereas multiparous females were middle-aged or old. Reproductive status was unknown for 10 mothers because they entered the study as adults with elongated nipples, indicating prior reproduction, but without information allowing us to distinguish primiparous from multiparous individuals. Eight of these females were categorised as middle-aged, one as young, and one as old. Although parity is conceptually a direct proxy for maternal experience, using parity would have reduced our sample size, whereas age category allowed us to retain all dyads and capture both parity status and cumulative maternal experience.

Four of the 49 mothers transferred to another unit or party during the study period. In two cases, the infant did not follow the mother (at 16.44 and 11.64 months of infant age). In one case, the infant followed the mother (age at transfer: 13.80 months). In the remaining case, the mother transferred to an unknown male outside the study parties when the infant was 2.79 months old. She subsequently returned to another male of one of our study parties when the infant was 4.92 months old; the infant followed the mother during both transfers. Two mothers died before their infant was 18 months old (infant age at disappearance: 12.88 and 12.61 months). In one of these cases, the mother had already resided in a different unit from her infant for 30 days before her disappearance. Subsamples included in each analysis are reported in each analysis section. All statistical analyses were conducted in the R environment (version 4.5.2: R Core Team, 2022) in the RStudio interface (version 2025.09.2.418: RStudio Team, 2022).

#### Behavioural data collection

During 2017-2021, we conducted 2361 focal protocols on 80 infants (45 females) using the focal-animal sampling method (Altmann, 1974). The observations lasted 15 minutes (median; IQR: 13.22-15.25 minutes). We recorded all approaches and leaves by the mother and infant (within 1 m) as well as the overall time an infant spent within 1 m of the mother. We also recorded the durations of nipple contact, contact-sit, maternal grooming of the infant, and carry (Dal Pesco & Fischer, 2022). Contact-sit, grooming, and carry were mutually exclusive. Nipple contact could co-occur with carry, contact-sit, and grooming. We additionally recorded rejection of nipple contact by the mother, rejection of carrying by the mother, and all aggression against the infant by the mother (e.g., mock bite, push away, threat) as events. Because of their rare occurrence, we combined all rejection behaviours into a single compound ‘rejection’ measure. Furthermore, we included protective behaviours (e.g., retrieving and restraining) as events. Data collection on rejection and protection behaviours began only in January 2019. All nine maternal behaviours listed above were used in our assessment of inter-individual variation in maternal style (Fig. 1). The detailed ethogram for maternal behaviours is provided in Tables S2.1 and S2.2.

**Figure 1.**
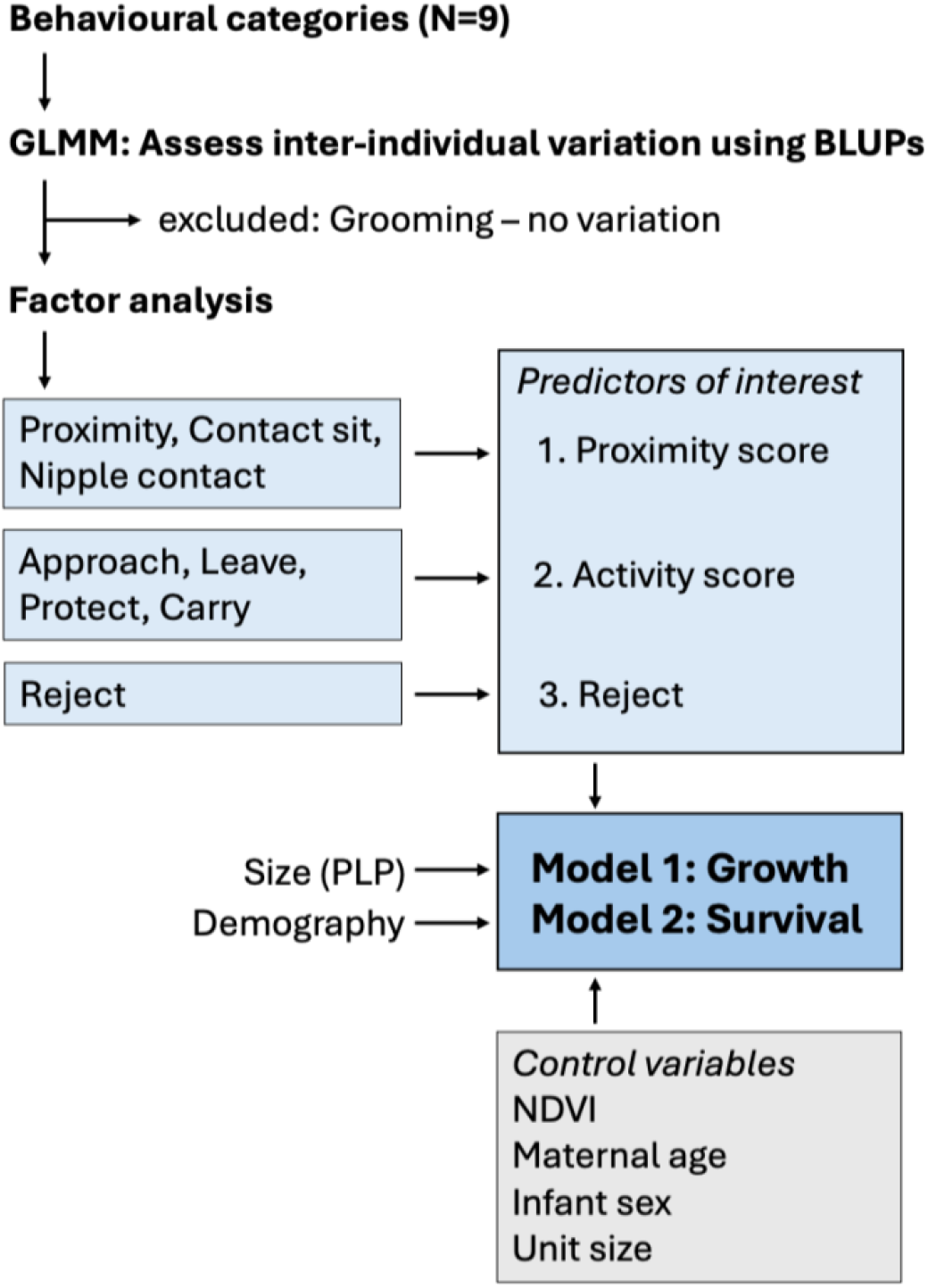
Workflow. We first screened the predictor variables for variation between mothers and infants using the estimated standard deviation of the BLUPs. We then used a principal components analysis to reduce the number of variables used to estimate variation in maternal style, yielding two factors and one separate variable (‘reject’). Lastly, we fitted the growth and survival models using the three resulting variables as predictors of interest, along with several control variables.

#### Parallel Laser Photogrammetry

We used Parallel Laser Photogrammetry (PLP) to determine infant growth over time. PLP is a non-invasive method that combines digital photography with parallel lasers to measure body size in wild animals (Anzà et al., 2022; Bergeron, 2007; Durban & Parsons, 2006; Galbany et al., 2017). We used a three-laser system in which the horizontal and vertical lasers were separated by 20 mm to measure the length of the lower arm in infants. We took pictures of each infant monthly from April 2019 to August 2021, for a total of 52 infants. Detailed explanations of the PLP in our study are provided in the supplement section S3.

We analysed a total of 1878 pictures (31 median number of images per individual; IQR: 12-57.75) using ImageJ version 1.53k, an open-source software package for scientific image processing and analysis (Schneider et al., 2012). Three researchers, trained in body landmark identification and length assessment, scored the images. The ICC(A,3) for assessing the inter-rater reliability indicated an excellent reliability (N pictures = 243, raters = 3, 0.993 < ICC < 0.995, F_(242,466)_ = 172, P < 0.001). For several infants, we measured arm lengths on the same day (*N* = 273 infant-day combinations with a maximum of 14 measures per day, including both arms). For these, we evaluated the range of arm-length measures obtained for the same infant on the same date. The average error (8.0 mm) was equivalent to 2 months of growth, with a maximum error (24.2 mm) equivalent to 8 months of development and a minimum error (0.2 mm) equivalent to ca. two days of growth.

#### Age-corrected determination of maternal style

To determine inter-individual differences in maternal style, we aggregated observations by infant and month, yielding a sample of 582 infant-month data points from *N* = 80 infants (45 females) born to 49 mothers. Since behavioural measures were aggregated by month, while infant age is a continuous measure, we first needed a monthly value of infant age to use in our analyses. To do this, we calculated an average age in days, weighted by the distribution of focal observations within the month. Maternal style for protect/reject behaviours could only be assessed from 2019 and was based on a sample of 425 infant-month data points from *N* = 64 infants (33 females) born to 45 mothers.

To describe mother-infant relationships, we first fitted separate Generalized Linear Mixed Models (GLMMs, Baayen, 2008), one for each of the nine maternal behaviours, with infant age (z-transformed) as a fixed effect, and mother ID and infant ID as random intercepts. These random intercepts allowed us to account for differences in maternal style originating from the different mothers included in the study, as well as potential variation in maternal style of the same mother toward her multiple infants. Infant age was included as a random slope within both mother ID and infant ID to account for variation in the effect of age on mothering style between mothers and infants (Barr et al., 2013; Schielzeth & Forstmeier, 2009). Behavioural states (grooming, contact-sit, nipple contact, carry, and proximity) were analysed using beta GLMMs fitted with the ‘glmmTMB’ function of the package ‘glmmTMB’ (version 1.1.13; Brooks et al., 2017). Count behaviours were analysed using Poisson (maternal approaches, protections, and rejections) or negative binomial (maternal leaves) GLMMs fitted with the ‘glmer’ and ‘glmer.nb’ functions of the R package ‘lme4’ (version 1.1-37; Bates et al., 2015). All count models included an offset term for observation effort (log transformed).

All models, except the ‘maternal leave’ model, were fitted without correlations between random intercepts and slopes because convergence problems occurred when they were included or they were estimated to be close to -1 or 1. For similar reasons, in the ‘maternal leave’ model, we kept correlations among random intercepts and random slopes only for age within mother ID, not for age within infant ID. We fitted models for maternal protection and grooming with an additional observation-level random effect to mitigate overdispersion. Across all nine models, overdispersion was detected only for grooming (dispersion parameter = 2.46), which we later excluded due to negligible variation in maternal style (see below). Dispersion parameters for the remaining models ranged from 0.45 to 1.10.

From the GLMMs, we obtained Best Linear Unbiased Predictors (BLUPs; Baayen, 2008). The BLUPs are the estimated deviations of intercepts and slopes from the common average for each level of the grouping factors mother ID and infant ID (Mundry et al., 2023). For each model, we first inspected the estimated standard deviations of the random intercept effects for mother ID and infant ID, but not those for random slopes, as random intercept BLUPs provide information about deviations of individual mothers’ maternal styles from that of the “average mother”. When both standard deviations were below 0.1, indicating little variation in maternal style among mothers or infants, we did not consider the results of the respective model in further analyses. We excluded grooming from further analysis because there was low variation between mothers and infants, while we retained the other eight variables (Fig. 1). In particular, BLUP assessment showed variation in the random intercepts corrected for infant age (SD > 0.1) between mothers for the variables carry, maternal leaves, protect, and reject. Maternal rejections also varied among infants, as did contact-sit, proximity, nipple contact, and maternal approaches. Population-average behaviours during the infancy period (0-18 months) are presented in Table S4.

To consider potential correlations among the behavioural variables and avoid the pitfalls of multiple testing, we conducted a principal components analysis (PCA) on the eight maternal behaviours that showed sufficient variation between mothers and infants (correlations between the eight variables are shown in Fig. S5). Because our models included random slopes of infant age within mother ID and infant ID the estimated variation associated with the random intercepts depended on the value of infant age. Since infant age was mean-centred in all models, random intercept variation was conditional on the average infant age in the dataset, which itself was influenced by heterogeneity in the longitudinal sampling structure (e.g., interruptions in data collection and unequal observation periods across infants). To obtain estimates that better reflected behavioural variation across development, we first generated infant specific fitted trajectories of maternal behaviour across the observed infant age range using the fixed effects estimates together with mother- and infant-specific deviations (i.e., Best Linear Unbiased Predictors; BLUPs; Baayen, 2008). We then quantified, for each mother-infant dyad, the average deviation between these dyad-specific fitted values and the fitted values expected for an average mother with an average infant (based solely on the fixed effects components; see also Fig. SI 25 in Mundry et al., 2023). These deviation estimates were subsequently used in our PCA. As we wanted to include all behaviours together, and deviation estimates for the protection and rejection models were only available for a subset of the data, our PCA was based on the subsample of 64 infants for whom all dyadic deviation estimates were available.

We settled on a two-factor solution and varimax rotation. The first factor had an Eigenvalue of 2.27 and explained 32.5% of the total variance. The highest loadings on the first factor (hereafter ‘proximity score’) were 0.858 for contact-sit, 0.828 for proximity, and 0.819 for nipple contact (Table S5). The second factor had an Eigenvalue of 1.27 and explained 18.1% of the total variance. The highest loadings on the second factor (hereafter ‘activity score’) were 0.704 for carry, 0.585 for maternal protection, 0.546 for maternal approach, , and 0.323 for maternal leave (Table S5). Maternal rejections did not load highly on either factor and was excluded from the final PCA. We therefore retained it as a third dimension of maternal style. In summary, the proximity score, activity score, and maternal rejection constitute three separate dimensions of maternal style in our dataset and were used as predictors in our main analysis, in which we examined the relationship between maternal style and infant growth and survival (Fig. 1).

#### Age-corrected determination of infant growth

We expressed infant growth as its deviation from the average growth of infants of the same age. To this end, we fitted a non-linear mixed-effects model, in which the response was arm length, and the principal predictor was infant age. We parametrized the link between size and infant age as follows: arm length = c + a · b^age^, where *c*> 0, *a* < 0, and *b* ranges from 0 to 1. Because 0 < b < 1, the term b^age^ decreases with increasing age; constraining a < 0 therefore ensures a monotonically increasing growth curve that asymptotically approaches c, consistent with biologically realistic growth. In the model, we let the coefficients *c*, *a*, and *b* vary between infants. We modelled them as fixed and random effects, whereby the latter implies that they are assumed to originate from a normal distribution with a standard deviation to be estimated. The model eventually provides estimates of *c*, *a*, and *b*, and how they vary across infants. This model was implemented in a Bayesian framework using the ‘brm’ function from the ‘brms’ package (Bürkner, 2017, 2018, 2021).

Next, we extracted the individual-specific infant growth parameters from the fitted model. We then determined the instantaneous fitted growth (i.e., the derivative of the fitted model) for each infant on each day between its first and last arm length measurement. Then we subtracted the population-level growth on the same day from each of these values. Finally, we averaged the deviations. These values, which are the slope differences, indicate how fast an infant grew relative to an average infant of the same age (i.e., relative infant growth). We then used the slope differences as a response in our growth models. We used infant growth rather than infant size because growth reflects changes in size over time. Differences in size between individuals of the same age may be due to genetic factors rather than to the conditions affecting infant development. Assessing individual growth trajectories in comparison to the average population growth trajectory for infants of the same age is a more accurate indicator of infant development.

#### Modelling the effect of maternal style on growth and survival

Of the 64 infants included in the principal components analysis to determine inter-individual differences in maternal style, one was excluded from both the growth and the survival models because full information on unit size, one of our control predictors, was unavailable due to uncertainty about its unit association. Thus, we included 63 infants (32 females) in the survival analysis. Note that six infants were not seen again after the COVID data gap and were therefore treated as censored data (i.e., alive until the last date seen before the COVID break). This sample included 44 mothers. Fourteen infants had young adult mothers, 41 had middle-aged mothers, and eight had old mothers.

The growth model included a smaller subset of infants because growth information was collected only from 2019 onwards. We included 52 infants for whom we had PLP pictures for the assessment of relative infant growth (Fig. 1). Of those, two had to be excluded from the growth model: one due to uncertain unit association (see above), the other because the infant died shortly after birth and no behavioural data could be collected. Therefore, for the growth model, we included 50 infants (25 females) born to 35 mothers. Nine infants had young adult mothers, 34 had middle-aged mothers, and seven had old mothers.

To analyse the link between maternal style and infant growth, we fitted a Linear Mixed Effects Model (LMM) using the ‘lmer’ function of the R package ‘lme4’ (version 1.1-37; Bates et al., 2015). For the link between maternal style and infant survival, we fitted a Cox proportional hazards model using the ‘coxme’ function of the package ‘coxme’ (version 2.2-22; Therneau, 2022). Mother ID and unit ID (i.e., the unit with which the infant was associated for most of its observed presence – from birth/first sighting to 18 months/last sighting) were included as random intercepts. No random slopes were theoretically identifiable. We used maternal age, infant sex, unit size, and the Normalized Difference Vegetation Index (as a proxy for ecological conditions; hereafter NDVI) as fixed-effect control factors (Fig. 1). Maternal age was defined as the mother’s age category on the day of each infant’s birth. During the study period, five unidentified females transferred to our study parties with an infant; for these cases, maternal age was assigned on their first observation day using our age category definitions (Dal Pesco & Fischer, 2022). Unit size for each infant was calculated as the weighted average number of unit co-residents (i.e., the primary male and all associated adult and subadult females) during the infant observed period. If an infant changed units during the period considered, we assigned it to the unit it belonged to for the longest period.

We obtained the NDVI values from the Copernicus Global Land Service (CGLS) (https://land.copernicus.eu/global/), which is part of the European Commission’s Earth Observation programme. See supplementary section S6 for details regarding NDVI extraction and processing. To control for the fact that we were forced to use two different NDVI versions, we created an additional indicator variable indicating whether the NDVI value for a given day was based on version 1 or version 2. Since some of the daily NDVI values were interpolated between NDVI versions 1 and 2, the indicator variable denoted the relative contribution of NDVI version 2 to a daily NDVI value. We deemed this step necessary because the two NDVI versions were not entirely comparable. NDVI was used as a fixed-effect control, together with its interaction with the indicator variable denoting the NDVI version.

Before fitting the two models, we inspected whether the distribution of the response variable (for the LMM) and all quantitative predictors was roughly symmetrical. Maternal rejections were square-root transformed in both models to improve symmetry; to ensure all values were positive, we added 1 in the growth model and subtracted the minimum value in the survival model. To ease model convergence and the comparison of estimates, we z-transformed all covariates to a mean of 0 and a standard deviation of 1 before fitting each model (Schielzeth, 2010). Before inference, we performed several diagnostic validations. Residuals showed no deviations from the assumptions of normality and homoscedasticity, as assessed by visually inspecting the residuals’ QQ-plot (Field, 2005) and a scatterplot of residuals versus fitted values (Quinn & Keough, 2002). Model stability was evaluated by comparing estimates obtained when running the models with the levels of the random effects excluded one at a time with those obtained for the complete data set (Nieuwenhuis et al., 2012). Both models were of acceptable stability.

As a general test of the statistical significance of maternal style and to avoid “cryptic multiple testing” (Forstmeier & Schielzeth, 2011), we conducted a full-null model comparison, whereby the null model lacked the three fixed effect predictors of maternal style (proximity factor, activity factor, and reject), but aside from that, was identical to the full model. We used a likelihood ratio test to compare the full and null models (Dobson, 2002). We employed the Satterthwaite approximation to test for the effects of the individual fixed effects in the infant growth LMM (Luke, 2017). To this end, we refitted the models with restricted maximum likelihood. For the survival analysis, we assessed the significance of the individual fixed effects by dropping them from the model one at a time (R function “drop1”). We obtained bootstrapped confidence intervals for model estimates and fitted values using the function ‘bootMer’ in the package ‘lme4’; we also used the function ‘simulate’ in the package ‘glmmTMB’ to obtain bootstrapped confidence intervals for duration behaviours included in Fig. 2. We determined degrees of freedom and *P*-values for the LMMs using the function ‘lmerTest’ of the package ‘lmerTest’ (Kuznetsova et al., 2017).

**Figure 2.**
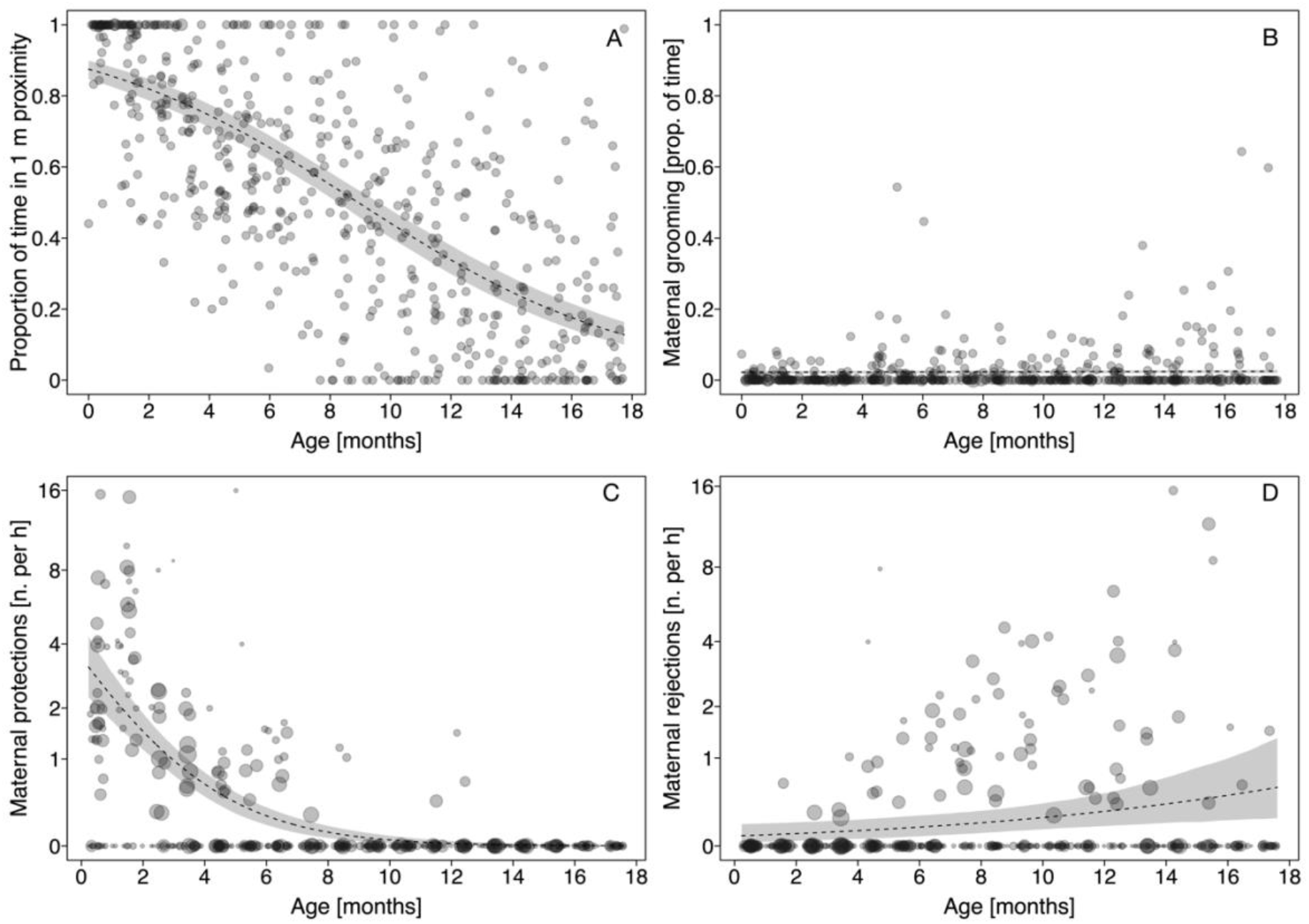
Raw data and estimates (mean and confidence intervals) for A: proportion of time in 1-m proximity; B: proportion of time the mother groomed the infant; C: number of maternal protections per hour. D: number of maternal rejections per hour.

#### Exploratory analysis of maternal style

We fitted two separate exploratory analyses, as per a reviewer’s suggestion. To test whether maternal age and infant sex explain variation in maternal style, we fitted separate Generalized Linear Mixed Models (GLMMs, Baayen, 2008), one for each of the eight maternal behaviours that varied between mothers and infants (i.e., excluding grooming). The model structures were the same as described above, except that the predictors maternal age and infant sex were added as fixed effects to the model as the predictors of interest. We used a likelihood ratio test to assess the significance of our full models relative to null models comprising all control predictors and the random effect structure, but lacking the two predictors of interest (Forstmeier & Schielzeth, 2011). We used the R function “drop1” to assess the significance of the individual fixed effects by dropping them from the model one at a time.

To explore the repeatability of maternal style across infants of the same mother, we created plots with the predicted values for each mother-infant dyad for mothers with multiple offspring (N=19), combining fixed effects with mother-level random effects (BLUPs) across the observed age range. We did this for each of the observed maternal behaviours that showed variation between mothers and infants (i.e., excluding grooming). We estimated repeatability using adjusted intraclass correlation coefficients (ICC; Nakagawa et al., 2017). Because our models included random slopes of infant age within mother identity, repeatability could not always be directly derived from the fitted models (see supplementary material S8). We therefore used two complementary approaches depending on model results: for models with negligible mother-specific age slopes, ICCs were calculated directly from the fitted models, whereas for models with non-negligible random slopes, we estimated mother-specific deviations from the population-level trajectories across infant age and quantified repeatability from these estimates using linear mixed-effects models. Further details are provided in the Supplementary Materials (S8).

## Results

### Maternal behaviour during development

In the first month of life, infants were mainly in close proximity to their mothers (Fig. 2A); time spent in close proximity declined with infant age, averaging 20% of the observation time at 18 months. Grooming did not vary with age and averaged around 2.2% of the observation time (Fig. 2B). Mothers showed high rates of protection behaviours in the first two months of the infants’ lives, with a decreasing rate until about 8 months of age, when they largely ceased to protect their offspring (Fig. 2C). With increasing infant age, maternal rejection became more frequent (Fig. 2D).

As shown by the PCA, maternal behaviours showed substantial correlations (Fig. S5). In particular, contact, proximity, and nipple contact were strongly positively correlated. Approach and leave behaviours, and carry and protect showed modest positive correlations. In contrast, rejection behaviours did not correlate with the other maternal behaviours.

We visualised maternal style across infants by plotting predicted values for each mother with multiple infants (N= 19), highlighting each infant, all other mother-infant combinations, and the population average. Representative examples of maternal consistency in proximity, nipple contact, and maternal rejection for mothers with 4 infants (N = 3) are shown in Fig. 3; plots for all 19 mothers with multiple infants are available in the supplementary materials (Fig. S7.1, S7.2, S7.3). Overall, maternal behaviours showed substantial variation both within and between mothers. Repeatability estimates assessed via intraclass correlation coefficients (ICC) were generally very low across maternal behaviours, indicating little consistency in maternal style across offspring within mothers. Only maternal rejection behaviour showed moderate repeatability (ICC = 0.51). In contrast, all other behaviours showed very low repeatability: the second highest ICC was 0.02 for carrying, whereas all remaining ICCs were below 0.01.

**Figure 3.**
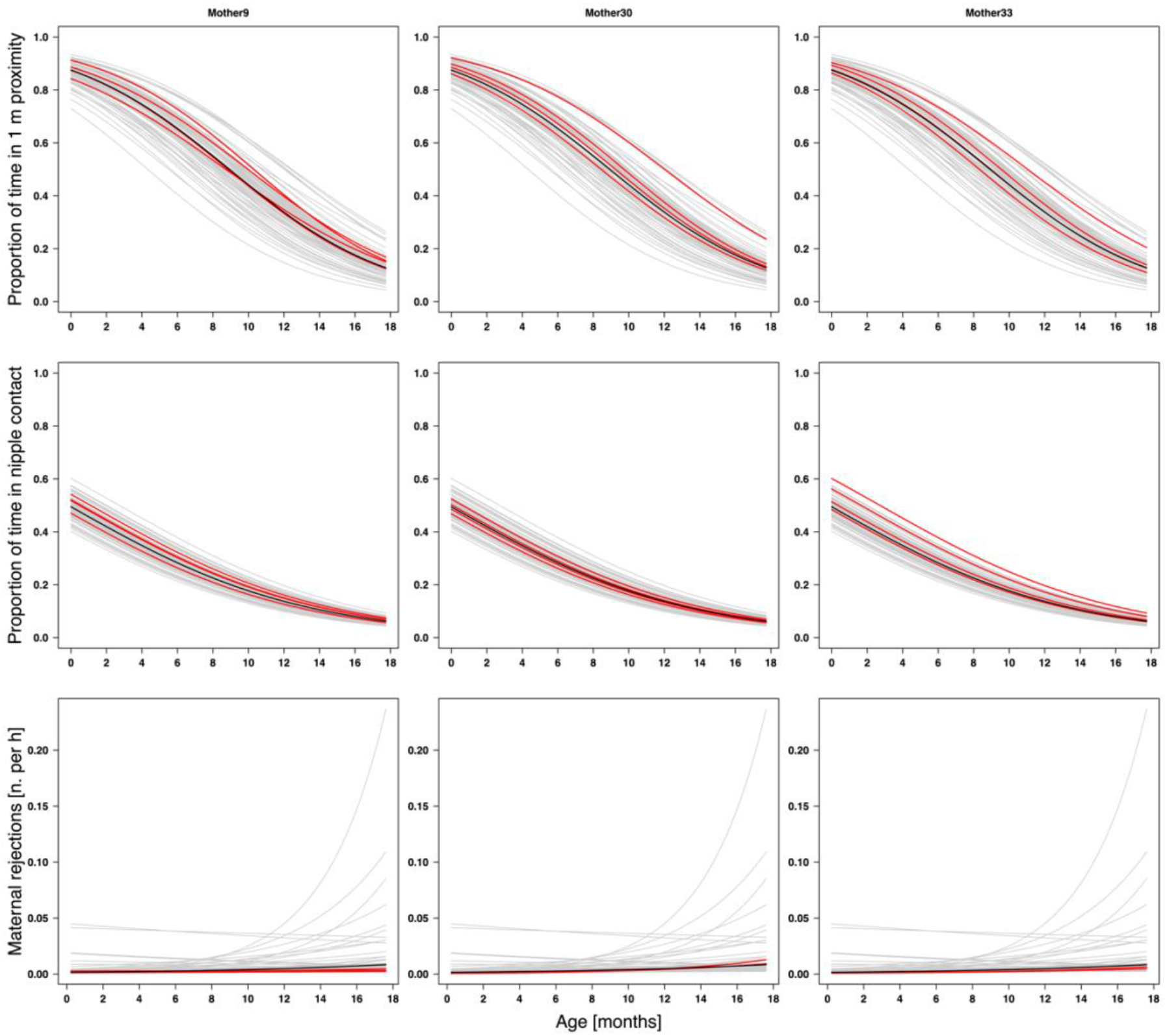
Predicted maternal behaviour across age for three representative mothers with four infants each for proportion of time in 1-m proximity (top panels), proportion of time in nipple contact (middle panels), and number of maternal rejections per hour (bottom panels). Each plot highlights a different mother with multiple infants, allowing visual assessment of maternal consistency and variation in each mother’s style relative to the other mothers and the population mean. Red lines show predicted trajectories for each mother and her four infants, grey lines represent all other mother-infant combinations, and black lines represent the population means.

Maternal age and infant sex did not explain variation in maternal style. For all eight maternal behaviours that showed variation between mothers and infants (i.e., excluding grooming), the full model including maternal age and infant sex did not explain significantly more variance than the null model (range of P-values = 0.08-0.739; see table S9 for each model results).

### Infant growth and survival

Of the 50 infants for which we had growth data (Fig. 4), 29 grew faster than the population average, and 21 grew more slowly. Infant death occurred similarly in the group of infants that grew faster than the average population (four infants) and in the group that grew more slowly (three infants). Due to the small sample size, we refrained from further statistical analysis. We extracted the estimated forearm length at 30 days to assess differences between males and females. For both sexes, the estimated forearm length at an age of 30 days was 78 mm. Similarly, the estimated forearm length at 30 days was 78 mm for both infants who survived and those who died.

**Figure 4.**
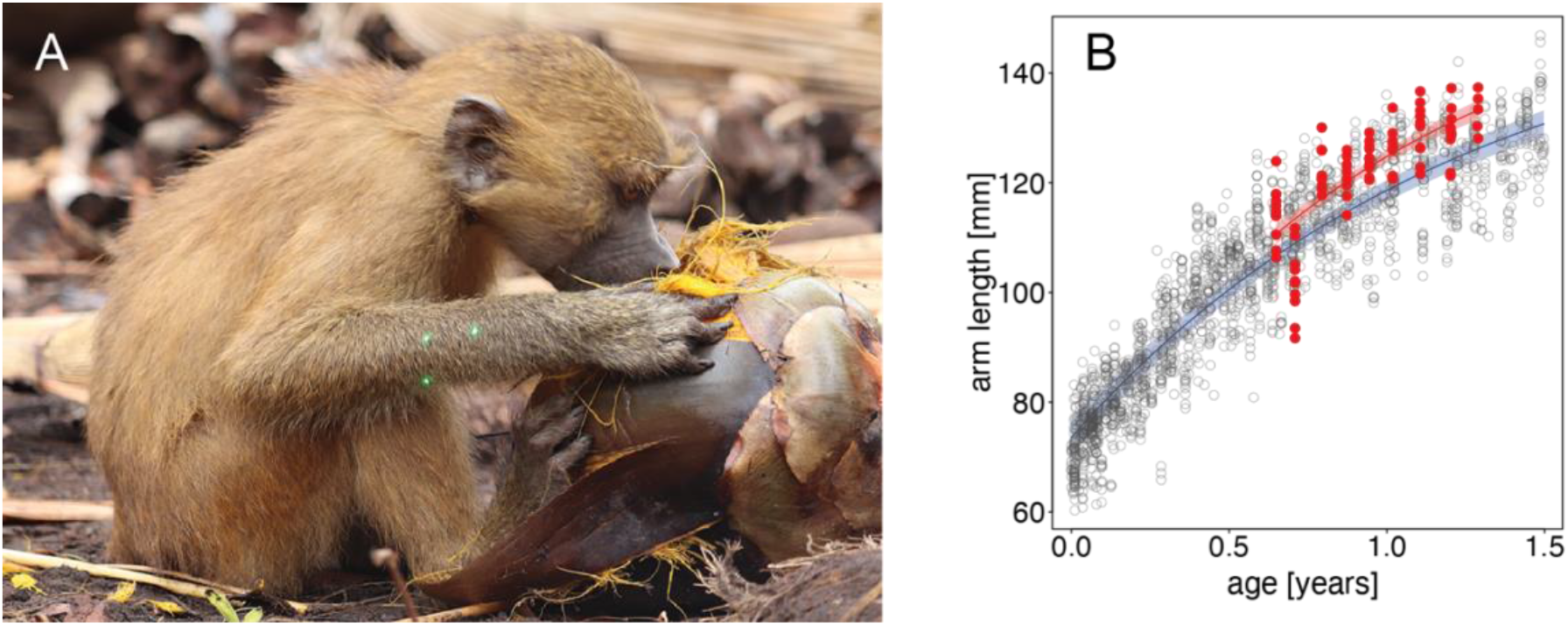
A. Measurement of arm length using Parallel Laser Photogrammetry of subject PIF. Note the three green spots on the infant’s forearm. B. Growth curve of the forearm length. Grey circles represent the arm-length measurements (N=1878) for all the infants of the population. The blue line represents the fitted model for the population average, and the red line depicts the fitted model for subject PIF. The shaded areas in blue and in red represent the 95% credible intervals of the fitted model for the average population and the subject PIF, respectively.

### Infant survival

Of the 80 infants in our initial sample, 63 survived until 18 months of age. Of the 17 infants who died, mortality occurred throughout the infancy period, with infants dying from an age of 13 days to 16.6 months. The highest infant mortality occurred in the first three months of life, during which seven infants died. Three infants died before reaching six months, four between six and nine months, and just one between nine months and one year. Two infants died after the first year: one was 13 months old, and the other was 16 months old. Of the 63 infants who entered the survival model (after the subsets described in the method section), 51 survived for up to 18 months. Note that these mortality rates are based only on infants for whom behavioural data were available and may therefore be underestimated, as infants that died shortly after birth and were not observed as focal animals are not included.

### Link between maternal style, growth, and survival

The growth model revealed no apparent relationship between maternal style and infant growth (Fig. 5). The full model with the three test predictors did not explain the data better than the null model (full-null model comparison: χ^2^ = 3.39, *df* = 3, *P* = 0.335). None of the test or control (infant sex, maternal age, unit size, and NDVI) predictors showed a noticeable effect on infant growth (Table 1).

**Figure 5.**
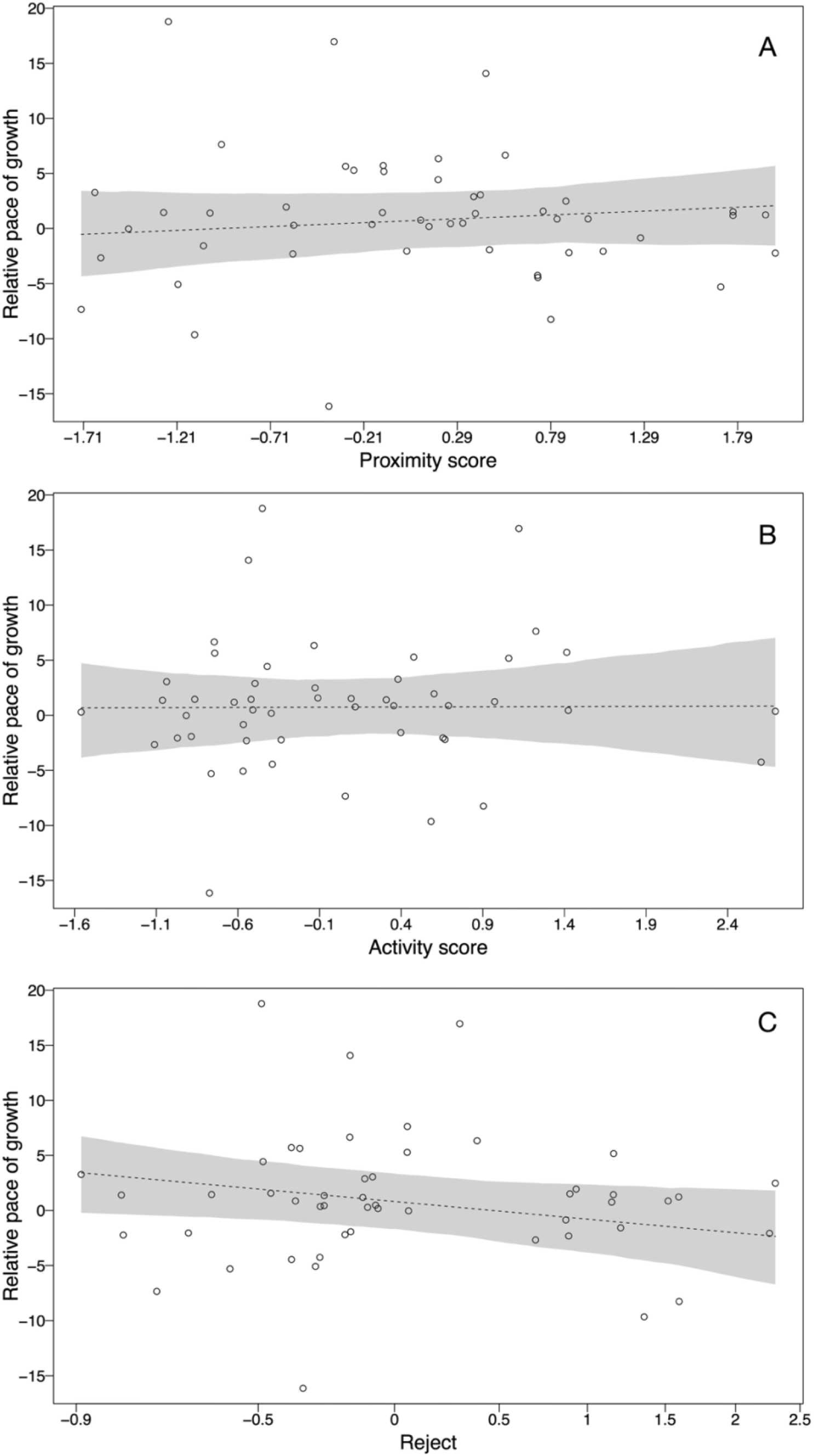
Relationships between the relative pace of growth of infants up to 18 months of Guinea baboons and (A) proximity score, (B) activity score, and (C) maternal rejections. Each circle represents an individual. The dashed line depicts the fitted model, and the shaded areas represent the bootstrapped 95% confidence intervals for all other terms in the model, assuming they are at their average.

**Table 1.**
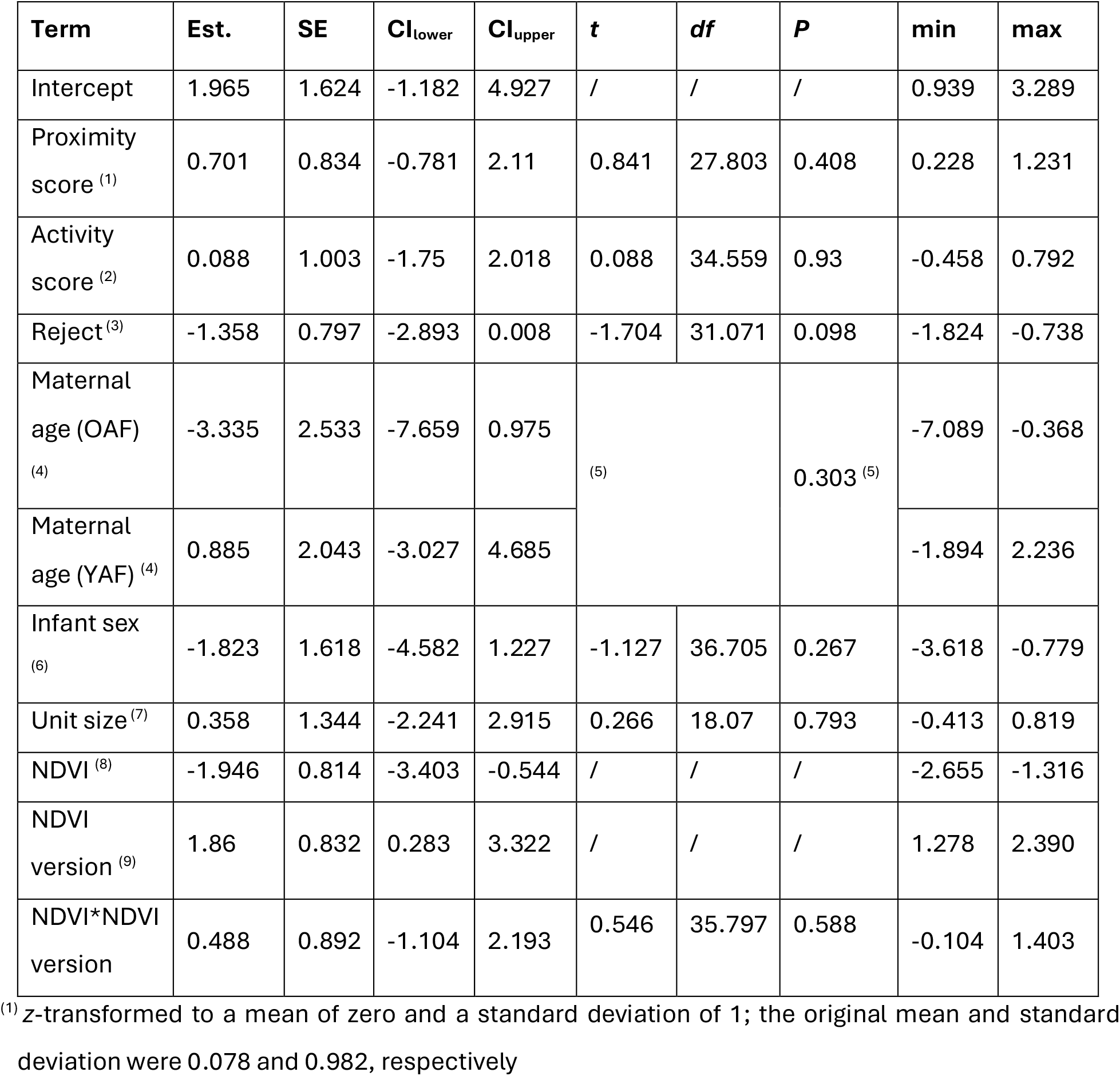
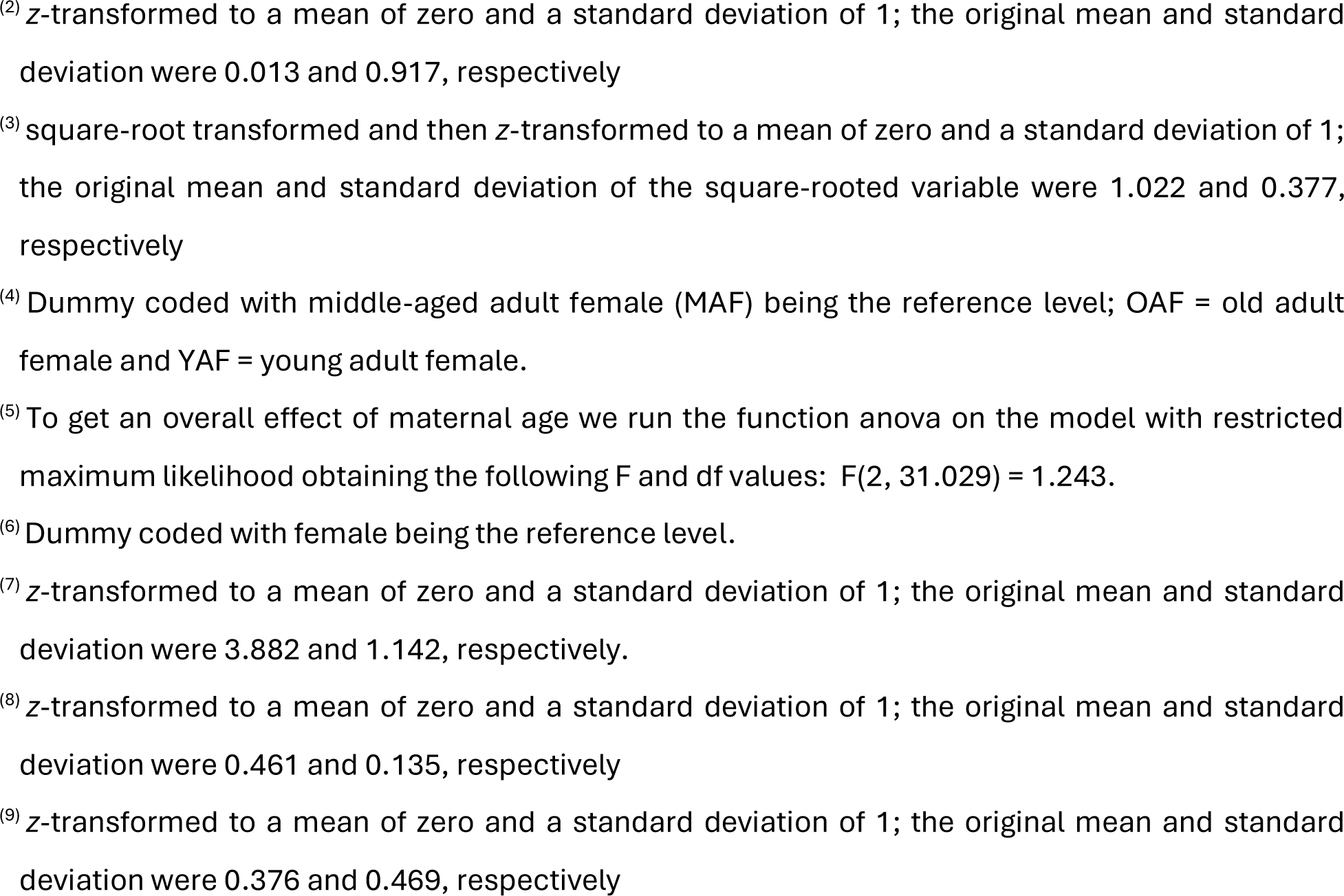
Results of the Linear Mixed Model (LMM) analysis of the influence of the test predictors of maternal style on infant growth (growth model). Control predictors are infant sex, maternal age, unit size, and NDVI values. Estimates are shown, along with standard errors (SE), 95% confidence intervals, significance tests, and minimum and maximum estimates from model stability analysis.

The survival model showed no apparent effect of maternal style on infant survival. The full model, which included the three test predictors, revealed no statistically significant relationships between the predictors and the dependent variables (full-null model comparison: χ^2^ = 0.292, *df* = 3, *P* = 0.962). None of the test or control (infant sex, maternal age, unit size, and NDVI) predictors showed a noticeable effect on infant survival (Table 2). Although male infants were almost twice as likely to die as female infants (10/35 vs. 7/44), mortality did not differ statistically between the sexes.

**Table 2.** Results of the Cox proportional hazard model analysing the influence of the test predictors of maternal style on infant survival. Control predictors are infant sex, maternal age, unit size, and NDVI values. Estimates are shown, along with standard errors (SE), significance tests, and minimum and maximum estimates from model stability analysis.

| Term | Estimate | SE | $\chi^2$ | df | P | min | max |
| --- | --- | --- | --- | --- | --- | --- | --- |
| Proximity score <sup>(1)</sup> | -0.124 | 0.343 | 0.13 | 1 | 0.718 | -0.271 | 0.129 |
| Activity score <sup>(2)</sup> | -0.097 | 0.312 | 0.1 | 1 | 0.752 | -0.543 | 0.381 |
| Reject <sup>(3)</sup> | 0.099 | 0.324 | 0.094 | 1 | 0.76 | -0.087 | 0.535 |
| Maternal age (OAF) <sup>(4)</sup> | 0.719 | 0.827 | 1.889 | 2 | 0.389 | 0.004 | 2.46 |
| Maternal age (YAF) <sup>(4)</sup> | -0.867 | 1.162 |  |  |  | -21.502 | -0.421 |
| Infant sex <sup>(5)</sup> | 0.816 | 0.658 | 1.605 | 1 | 0.205 | 0.407 | 1.542 |
| Unit size <sup>(6)</sup> | -0.106 | 0.381 | 0.078 | 1 | 0.78 | -0.557 | 0.316 |
| NDVI <sup>(7)</sup> | 0.221 | 0.424 | - | - | - | -0.589 | 0.822 |
| NDVI version <sup>(8)</sup> | 0.026 | 0.473 | - | - | - | -0.61 | 0.662 |
| NDVI*NDVI version | 0.975 | 0.469 | 1.881 | 1 | 0.17 | 0.179 | 1.994 |

<sup>(1)</sup> z-transformed to a mean of zero and a standard deviation of 1; the original mean and standard deviation were 0.04 and 0.889, respectively
<sup>(2)</sup> z-transformed to a mean of zero and a standard deviation of 1; the original mean and standard deviation were 0.014 and 0.84, respectively
<sup>(3)</sup> square-root transformed and then z-transformed to a mean of zero and a standard deviation of 1; the original mean and standard deviation of the square-rooted variable were 1.013 and 0.361, respectively
<sup>(4)</sup> Dummy coded with middle-aged adult female (MAF) being the reference level; OAF = old adult female and YAF = young adult female; the indicated significance test is an overall test of the effects of maternal age
<sup>(5)</sup> Dummy coded with female being the reference level
<sup>(6)</sup> z-transformed to a mean of zero and a standard deviation of 1; the original mean and standard deviation were 4.031 and 1.073, respectively.
<sup>(7)</sup> z-transformed to a mean of zero and a standard deviation of 1; the original mean and standard deviation were 0.529 and 0.118, respectively
<sup>(8)</sup> z-transformed to a mean of zero and a standard deviation of 1; the original mean and standard deviation were 0.433 and 0.455, respectively

### Qualitative assessment of protection and rejection behaviours

To facilitate comparisons with previous data on yellow baboons (Altmann, 1980), we additionally considered only those mother-infant dyads for which we had data on the first four months (i.e., cases recorded from the time we began to record protection and rejection behaviours; N = 15). Our qualitative assessment showed that two mother-infant dyads exhibited no protective behaviours at all, while in the remaining 13 dyads mothers protected their infants on average 2.06 times per hour (SD = 1.43) over the first four months. Among these 13 dyads, protective behaviours were observed at least once per month in six dyads, but not in the remaining seven. Of the 15 infants in this sample, 14 survived to 18 months of age. All infants for whom we had no records of protection in the first four months survived, while of the six infants with the highest rates of protection, only three survived.

## Discussion

The primary aim of this study was to characterise maternal styles in wild Guinea baboons and examine their effects on infant growth and survival up to 18 months. Although mothers varied in their social investment, we found no evidence that dimensions of maternal style, including the maternal contact score (time in contact-sit, 1-m proximity, and nipple contact), the maternal activity score (maternal approaches, leaves, protections, and carry durations), or maternal rejections, influenced infant growth or survival.

The finding that maternal style had no evident effects on infant growth or survival may be due to methodological or sample limitations, or it may reflect a genuine absence of an effect. We found that the PLP method for assessing growth was associated with significant measurement error. While the interobserver reliability in measuring the inter-laser distance, recognising body landmarks in the picture, and measuring arm length in pixels was excellent, the measurement error in the field was considerable, and possibly larger than any variation due to maternal care. In addition, our dataset was characterized by many subjects but significant heterogeneity. While we addressed heterogeneity by employing an analytical approach that controlled for infant age-related behavioural changes and data gaps when analysing ontogenetic processes (Mundry et al., 2023), our measure of maternal style may still be uncertain. In sum, the high uncertainty in both the predictor (maternal style) and one of the response variables (infant growth) may have contributed to our failure to detect effects on growth. In the survival analysis, however, the response variable was unambiguous.

It is also possible that maternal style truly does not affect infant growth and survival in our population. In ecological contexts in which food availability is high and feeding competition is limited, mothers may experience broadly similar energetic conditions (Clutton-Brock, 1991), potentially reducing both the need for and consequences of variation in maternal care. Consistent with this interpretation, our study population inhabits a habitat with a high carrying capacity and shows little overt food competition (Ohrndorf et al., 2025a). Under such conditions, maternal energetic state might not differ substantially, and the observed variation in maternal style may still provide sufficient care to ensure normal infant growth and survival. Yet, the observed variation in maternal style raises the question of which factors may contribute to these differences. We found no evidence that maternal age or infant sex were related to maternal style (see also Arbaiza-Bayona et al., 2026 who found no effect of infant sex in Assamese macaques, *Macaca assamensis*). The observed variation between mothers could, in theory, also be due to stable inter-individual differences (‘personalities’), but we found no evidence for intra-individual consistency in maternal style. Alternatively, this variation may reflect behavioural flexibility, with mothers adjusting their behaviour in response to environmental and social conditions to buffer potential adverse effects on their infants. If effective, such flexibility may also reduce differences in infant growth and survival, thereby contributing to the absence of detectable effects of maternal style. Beyond postnatal care, prenatal effects (e.g., Berghänel et al., 2016) may be more important than postnatal care in determining infant health and growth, although this also seems unlikely given the lack of overt food competition and little variation in GC levels among females (Goffe et al., 2025). In summary, the sources of variation in maternal behaviour remain unexplained, and this variation does not seem consequential for growth and survival in early development. We suspect that mortality is mainly related to disease or injuries in young infants and predation, once young become more independent. Whether infant disappearances are indeed due to predation is hard to determine, however, as key predators such as leopards, *Panthera pardus*, are mostly active at night, and predation by snakes or birds of prey would be difficult to detect.

Infant growth rate and survival were not related to infant sex in our sample. In a comparative analysis of neonatal body mass in 109 primate species, neonatal sexual dimorphism positively correlated with adult dimorphism (Smith & Leigh, 1998). Despite the striking differences in adult body mass among Guinea baboons (Fischer et al., 2017), we found no evidence of sex-related differences in growth or estimated size during infancy. Similarly, Anzà and colleagues (2022), using the PLP method in Assamese macaque infants, reported no differences in forearm length during the first year of life; sexual dimorphism in forearm length emerged only once males reached 5-7 years of age. The emergence of sexual dimorphism in Guinea baboon body size appears later in life; therefore, it occurs in the absence of detectable sex-biased maternal investment, arguing against early maternal allocation as a primary driver of sexual dimorphism.

To account for potential ecological conditions that may influence growth and survival in our sample of infants, we also included NDVI as a fixed control effect. Again, we found no apparent effect on infant growth and survival. However, we now know that NDVI is a poor proxy for food availability in our population because several key fruiting tree species produce ripe fruit at the peak of the dry season, when NDVI greenness is lowest (Ohrndorf, 2025). In a one-year-long study, food availability scores were also the highest during this period (Ohrndorf et al., 2025a). In hindsight, we would not include this variable again. Rainfall does not provide a better proxy for food availability than NDVI (Ohrndorf, 2025). Some fruit only ripen in the dry season, and the animals also rely on fallback foods such as herbaceous vegetation and invertebrates (Ohrndorf et al., 2025a).

Maternal age was not a prominent determinant of infant growth and survival. Although maternal age and maternal experience are conceptually distinct, they were closely linked in our dataset: all primiparous females were classified as young, while middle-aged and old females were multiparous. Our age categories reflect increasing cumulative reproductive experience rather than a binary contrast between first-time and experienced mothers. The absence of an age effect, therefore, suggests that neither early reproductive inexperience nor incremental experience gained across multiple successive births has consequences for infants in our population. This finding stands in stark contrast to observations for other primate species, where younger mothers often experience a higher infant mortality than older ones (e.g., four groups of habituated wild grey-cheeked mangabeys, *Lophocebus albigena*, in Uganda: Arlet et al., 2014; six out of seven studies from captive and wild primate populations across different species: Pusey, 2012). One possibility is that favourable ecological conditions, along with social support from unit members (i.e., the primary male and co-resident females) within a generally tolerant environment, may buffer inexperienced mothers from resource and skill-related constraints, allowing infants of young mothers to develop just as successfully as those of more experienced mothers.

Comparing maternal style across species is challenging due to differences in social organization, ethograms, and data-collection protocols. To illustrate these challenges, we compare our results in Guinea baboons with published data from yellow baboons in Amboseli. This species lives in multi-male-multi-female groups with female philopatry, resulting in societies organised around matrilines (Alberts, 2019 for a synthesis). At six months of age, mothers and infants were within 1 m of each other around 48% of the time in yellow baboons (Altmann, 1980), compared with 65% of the time in Guinea baboons. The high overall gregariousness of the Guinea baboons (Goffe et al., 2016) may account for this difference, but more comprehensive comparisons would be needed to test this conjecture.

Maternal carry behaviours also differ between these species. Shortly after birth, mothers in both species spent a similar proportion of time carrying infants: around 30% in yellow baboons (Altmann & Samuels, 1992) and 42% in Guinea baboons. By 6 months, carrying in yellow baboons decreased sharply to 5% and was nearly absent by eight months, whereas in Guinea baboons, it remained 24% at 6 months and persisted at 6% by 18 months. Disentangling whether these differences reflect maternal style or environmental conditions is difficult. For example, females that need to travel longer distances for foraging may carry their infants for longer periods of time. Reliable comparisons, therefore, require means to assess the influence of other variables (e.g., information on time spent moving) and, ideally, identical data collection protocols (Kalbitzer et al., 2015).

Altmann (1980) classified yellow baboon mothers as “laissez-faire” and “restrictive”, based on the time at which they ceased restraining their infants (≤ 1 month versus 1.5 months or older). Mothers in our sample whose infants reached at least an age of 4 months and whose infants were sampled throughout this period also showed marked variation. Applying Altmann’s classification to our Guinea baboon subsample, some mothers would be considered “laissez-faire”, providing relatively little overt protection, whereas those providing high levels of protection could be classified as “restrictive”. Protection, however, is not solely driven by maternal style; it can also be considered as a response to the infant’s behaviour. Infants who continue to cling to their mothers do not prompt maternal intervention, while more active infants may more readily evoke protective behaviour in mothers. Thus, infants are not only passive recipients of maternal behaviour but actively shape the relationship (Fairbanks & Hinde, 2013). Likewise, when mothers are more rejecting, infants may respond by increasing their effort to maintain proximity (Fairbanks & Mcguire, 1995). Future studies should consider the interplay between infant behaviour and maternal responses in more detail, as it may be crucial for understanding variation in early social development.

There is a trade-off between dense sampling of a few infants or more sparse behavioural sampling of a relatively large number of infants, as in our study. Altmann (1980) initially sampled 12 yellow baboon infants. However, the yellow baboon population experienced high infant mortality, and only seven infants were present at the end of the twelve-month age window. In contrast, at twelve months of age, we still had 29 mother-infant dyads. A very dense sampling of a few infants allows solid inferences for these individuals. Paul (1984) observed seven infant Barbary macaques every other day, totalling 544 observation hours per infant over 2 years (except for one infant that died). Yet, in such a small sample, one or two extreme cases can distort the overall picture, and statistical inferences remain limited. In contrast, sparse sampling of many infants can better approximate the population average. Because limited information is available for any given individual, it becomes difficult to assess how strongly they deviate from the population mean. The structure of the data may have affected our ability to estimate maternal style variation and its contribution to infant growth and survival. Therefore, given our broad infant sample but limited data per mother–infant dyad, we are more confident about the average developmental trajectories of the infants than the variation between mothers. Although the comparison between species, notably Guinea and yellow baboons, must be taken with a pinch of salt, the observed differences appear to be in degree and not in kind. The observed mother-infant relationships follow a common pattern across baboon (and other catarrhine) species. We propose that differences between baboon species emerge only later in life, as part of physical and physiological maturation, ultimately leading to the pronounced differences in bonding patterns observed between baboon species (Fischer et al., 2019).

While numerous studies have investigated differences in maternal style across taxa (Georges & Guinet, 2000; Hill et al., 2007; Lee & Moss, 1986; Mann & Watson-Capps, 2005), fewer have also considered the consequences for growth and survival (see Altmann, 1980, for a small set of infants). Outside primates, the most direct tests of whether variation in maternal behaviour predicts offspring growth or survival are available in ungulates. In mountain goats, *Oreamnos americanus*, maternal care behaviours significantly predicted kid survival (Théoret-Gosselin et al., 2015), and in white-tailed deer, *Odocoileus virginianus*, both experimental and observational studies have linked the timing of visits to cached neonates and variation in lactation rates to fawn growth and survival under predation risk (Muthersbaugh et al., 2024; Therrien et al., 2008). By contrast, rodent models provide strong evidence for stable individual differences in maternal care, but the link to growth or survival outcomes has not been established (e.g., Champagne et al., 2003). Yet, none of these studies used the concept of maternal style, and the behaviours under investigation differed substantially from those observed in primate mothers and infants.

In summary, we observed variation in maternal style across a wide range of maternal behaviours, particularly regarding protection, rejection, carrying behaviour, and maternal approaches and leaves. Nevertheless, we found no link between variation in maternal style and infant growth and survival. Most likely, the observed variation in our study population falls within the species-typical reaction norm, such that moderate differences in maternal behaviour do not translate into measurable fitness consequences during early development. In other words, early development in this population may be relatively robust to naturally occurring variation in maternal behaviour. Only more drastic events, such as the loss of the mother or severe droughts (Tung et al., 2016), are likely to have fitness consequences during early development. Once we can consider the long-term effects of adverse events in early development, we can submit this conjecture to empirical testing.

## Acknowledgements

We thank the Diréction des Parcs Nationaux (DPN) and the Ministère de l’Environnment et de la Protéction de la Nature (MEPN) de la République du Sénégal for approval to conduct this study in the Parc National du Niokolo-Koba (PNNK). The support and cooperation of former and present park conservators Mallé Gueye, Amar Fall, Assane Ndoye, and Jacques Gomis are particularly appreciated. We are grateful to all the CRP Simenti staff and field assistants; in particular we thank the CRP Simenti park rangers El’Hadji Yankhoba Dansokho, Touradou Sonko, Vieux Biaye, Djibril Coly, Chérif Younousse Kéba Camara, and Amadou Bamba Diedhiou for their support in the field. We are thankful to Dominique Treschnak, Irene Gutiérrez Díez, Lisa Ohrndorf, Josefine Kalbitz, and Rachel Sassoon for their help and data collection in the field. We also thank Laura Camón, Lidia Jiménez, and Maren Decker for their assistance in PLP measuring. This research was supported by the Deutsche Forschungsgemeinschaft (DFG, German Research Foundation), Research Training Group 2070 „Understanding social relationships“, Grant/Award Number: 254142454 / GRK 2070.

## Author Contributions

Conceptualization: AAdD, FDP, JF; Investigation: AAdD; Methodology: AAdD, FDP, RM, JF; Software: FDP, RM; Formal analysis: AAdD, FDP, RM, JF; Data curation: AAdD, FDP, JF; Writing – Original Draft: AAdD; Writing – Review & Editing: AAdD, FDP, RM, JF; Visualization: AAdD, FDP, RM, JF; Funding acquisition: JF.

## Data availability statement

The data and code for the statistical analyses that support the findings will be made publicly available on OSF upon acceptance of the manuscript.

## Supplementary material

### S1. Behavioural data availability

**Figure S1.**
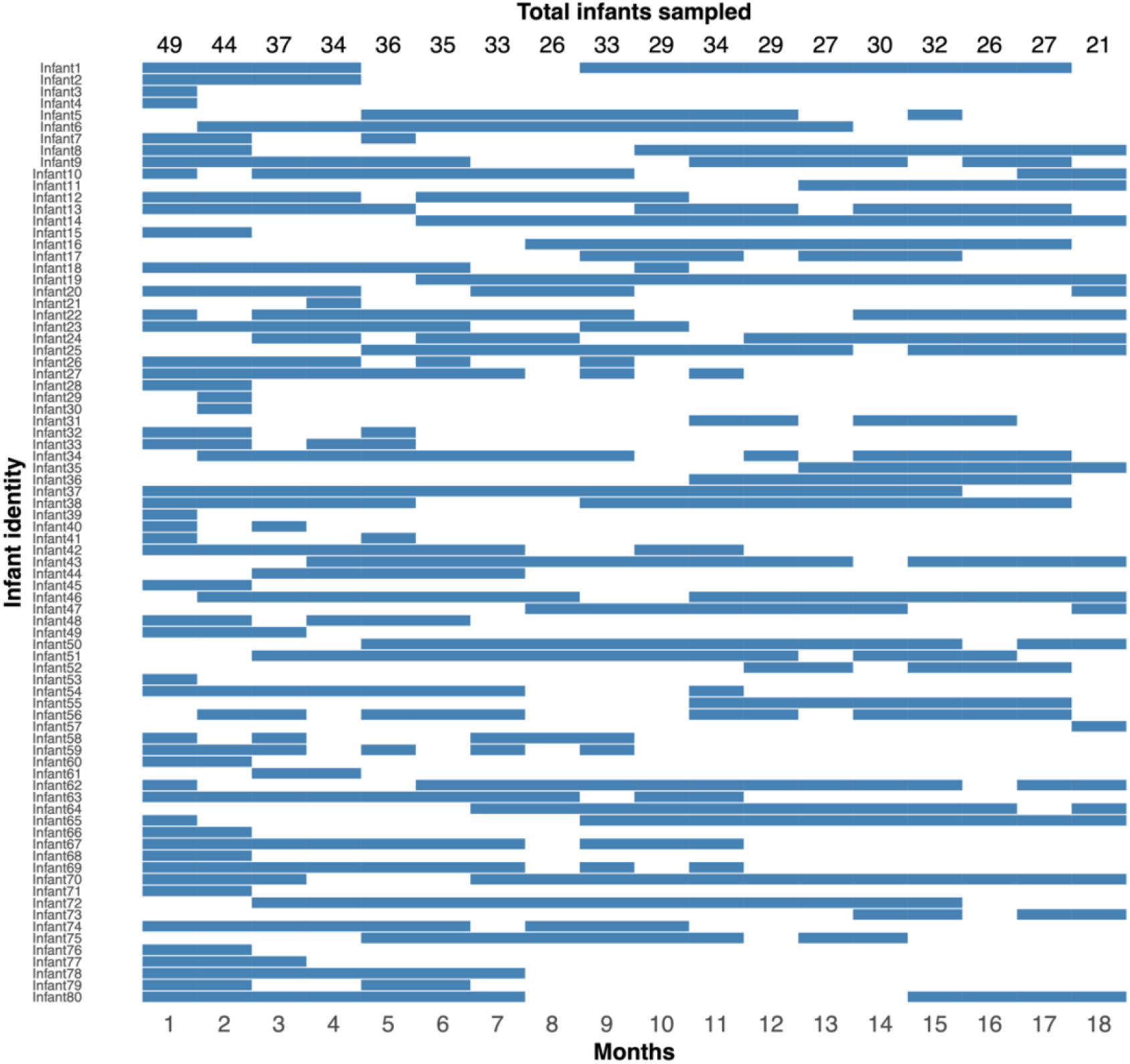
Overview of the sample sizes for behavioural data collection during the study period (April 2017 – December 2021). The blue cells depict monthly data availability for the 80 studied infants from birth until 18-months of age. On the top the total number of infants sampled per month is reported (mean= 32.3, range= 21-49). Data collection gaps are due to multiple reasons: infants were born before the study started (i.e., no data are available from birth), infants were born less than 18-months from the study period end (i.e., no data collection until 18-months of age), infants transferred into/out of our study parties (i.e., no data available before/after the transfer), study parties or individuals not found for a time period, COVID-19 data collection break (from April to November 2020)

### S2. Ethogram

**Table S2.1.** Ethogram of state behaviours between a mother and her infant that were considered for the FAs. Adapted from the CRP Simenti work manual (Dal Pesco & Fischer, 2022; version August 2022)

| Behaviour | Description |
| --- | --- |
| Maternal grooming | The mother moves her fingers and/or lips through the fur of her infant. A grooming bout is considered finished only once the mother has stopped grooming her infant for more than 5 s. Grooming, contact-sit, and carry are mutually exclusive. |
| Contact-sit | Mother and infant are sitting, lying, or standing within < 10 cm of one another, meaning that the individuals either touch skin-to-skin or are so near that their furs touch, with at least one individual sitting or lying down. This applies to the body, the hairs, and the different body parts (e.g., feet or hands) of the two individuals. However, if one individual is in contact with another one only with its tail, body contact is recorded only if it is the base of the tail that is in contact. A contact-sit bout is considered finished only once contact stopped for more than 5 s. Grooming, contact-sit, and carry are mutually exclusive. |
| Proximity | Total time that the mother and infant are within 1 m proximity during the observation time. This is calculated using information on:<br>- proximity at the beginning and end of the focal protocol<br>- detailed recordings of movements during the observation time (i.e., approaches into 1 m and leaves out of 1 m between the mother and infant) |
| Nipple contact | The infant has its mother's nipple in its mouth, either holding onto it or suckling. This behaviour is not mutually exclusive to contact sit, carry, and grooming. |
| Carry | The infant is carried by its mother, either ventrally or dorsally. If the mother stops walking but the infant is still holding on to the mother's fur with all four extremities, it is still carrying behaviour. However, when the mother is not walking and the immature is grabbing the mother's fur only with the forelimbs, this is not coded as carried, but as contact sit. Grooming, contact-sit, and carry are mutually exclusive. |

**Table S2.2.** Ethogram of event behaviours between a mother and her infant that were considered for the FAs. Adapted from the CRP Simenti work manual (Dal Pesco & Fischer, 2022; version August 2022)

| Behaviour | Description |
| --- | --- |
| Protection | Aggregate of two protective behaviours: retrieve and restrain. |
|  | *Retrieve*: the mother recovers the infant (including from other individuals) by grabbing or holding it with one or both arms impeding that the infant moves away from her. |
|  | *Restrain*: the mother stops the infant from moving away by grabbing and holding the infant. |
| Rejection | Aggregate of rejecting behaviours and aggressive behaviours directed from the mother to the infant. |
|  | *Rejection of nipple contact*: the mother avoids her infant from starting nipple contact. |
|  | *Rejection of carrying*: the mother prevents her infant from being carried. |
|  | *Mock-bite*: the mother bites her infant with moderate strength. |
|  | *Push away*: the mother presses or shoves her infant away from her, avoiding either contact or proximity between them. |
|  | *Threat*: the mother stares at the infant with lifted eyebrows or stands on all four limbs with her hands placed slightly in front of the body. The mother can also conduct a small but fast movement forward while all legs and arms remain on the ground, or one arm loses ground (ground slap). |
|  | *Physical fight*: The mother hits or bites or throws the infant on the ground. |
|  | *Submissive behaviour*: the infant displays submissive behaviour such as fear grin, crouch, or tail-raise towards the mother or via vocalisations such as keck or scream. |
| Maternal approaches | The mother decreases her distance to the infant to less than 1 m. |
| Maternal leaves | The mother increases her distance to the infant to more than 1 m. |

### S3. Parallel Laser Photogrammetry

The apparatus, or Parallel Laser Photogrammetry (PLP) system, consisted of a digital single-lens reflex (DSLR) Camera Canon EOS 200D and an objective Canon EFS 55-250mm, along with a laser box that contained three parallel lasers (laser class 2 520 nm 1mW DI520-1-3) (Fig. S3.1). We chose a three-laser system because a third laser creates a right isosceles triangle with the other two lasers, which allows to become aware and correct whenever the photographed object deviates from the perpendicular orientation of the camera-object axis, or to detect whether the photographed object has tilts on the surface (Anzà et al., 2022; Galbany et al., 2016). In both cases, the right angle of the triangle would be lost, as one of the laser dots would appear displaced (see Anzà et al., 2022). We used green lasers, as in daylight they are more visible than red lasers (Bergeron, 2007; Durban & Parsons, 2006). Three PLP systems were used during the duration of the study: PLP1, PLP2, and PLP3.

**Figure S3.1.**
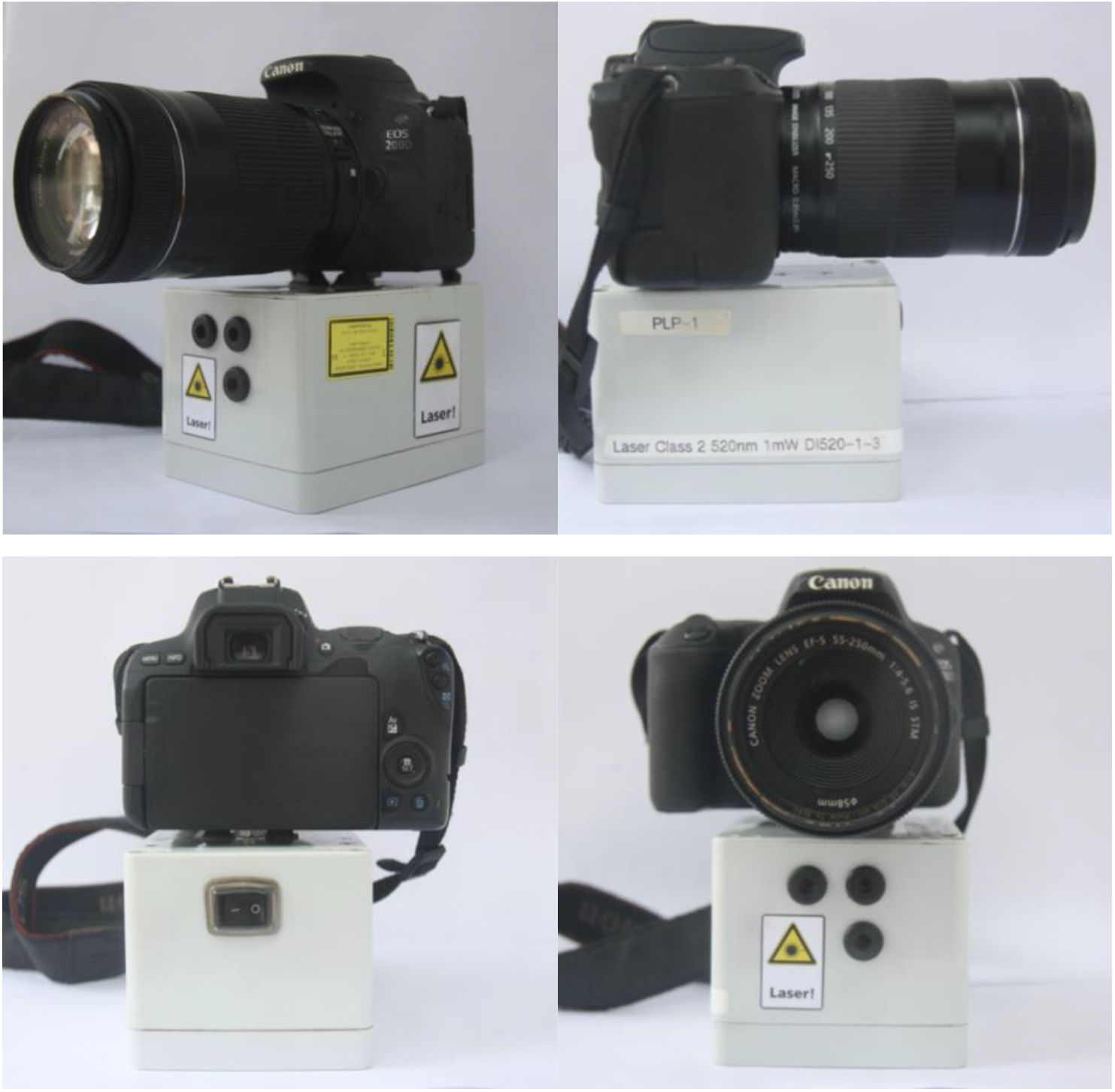
Apparatus for Parallel Laser Photogrammetry (PLP) used in the study, seen from different angles. The apparatus, also referred to as PLP system, consisted of a camera and a laser box with three lasers and a switch. Batteries were located inside the laser box. Horizontal and vertical lasers were separated by 20 millimetres

The number of pixels between the paired lasers projected on the surface of the body (inter-laser distance) that appear in the photo are used as a scale (Richardson et al., 2022) since a relationship can be made between inter-laser distance and the already-known inter-beam distance (see formula below). The scale is then used to convert the arm size in pixels (determined by the distance between body landmarks) into arm size in millimetres (see formula below). Unfortunately, due to calibration issues in the field, only the horizontal paired lasers could be used as a scale (see below for details about calibration issues).

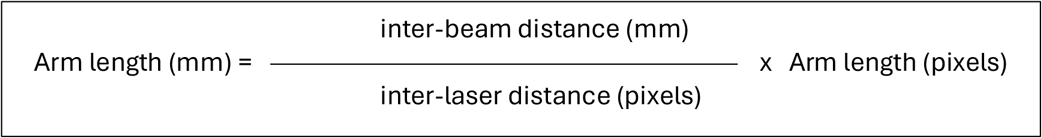

The individuals were photographed in their natural habitat, in lateral view. Observer distance was about 2 to 5m. To determine the length of the lower arm, we defined two landmark endpoints for our measurements (Fig. S3.2, A-B): the olecranon and the ulnar styloid process.

**Figure S3.2.**
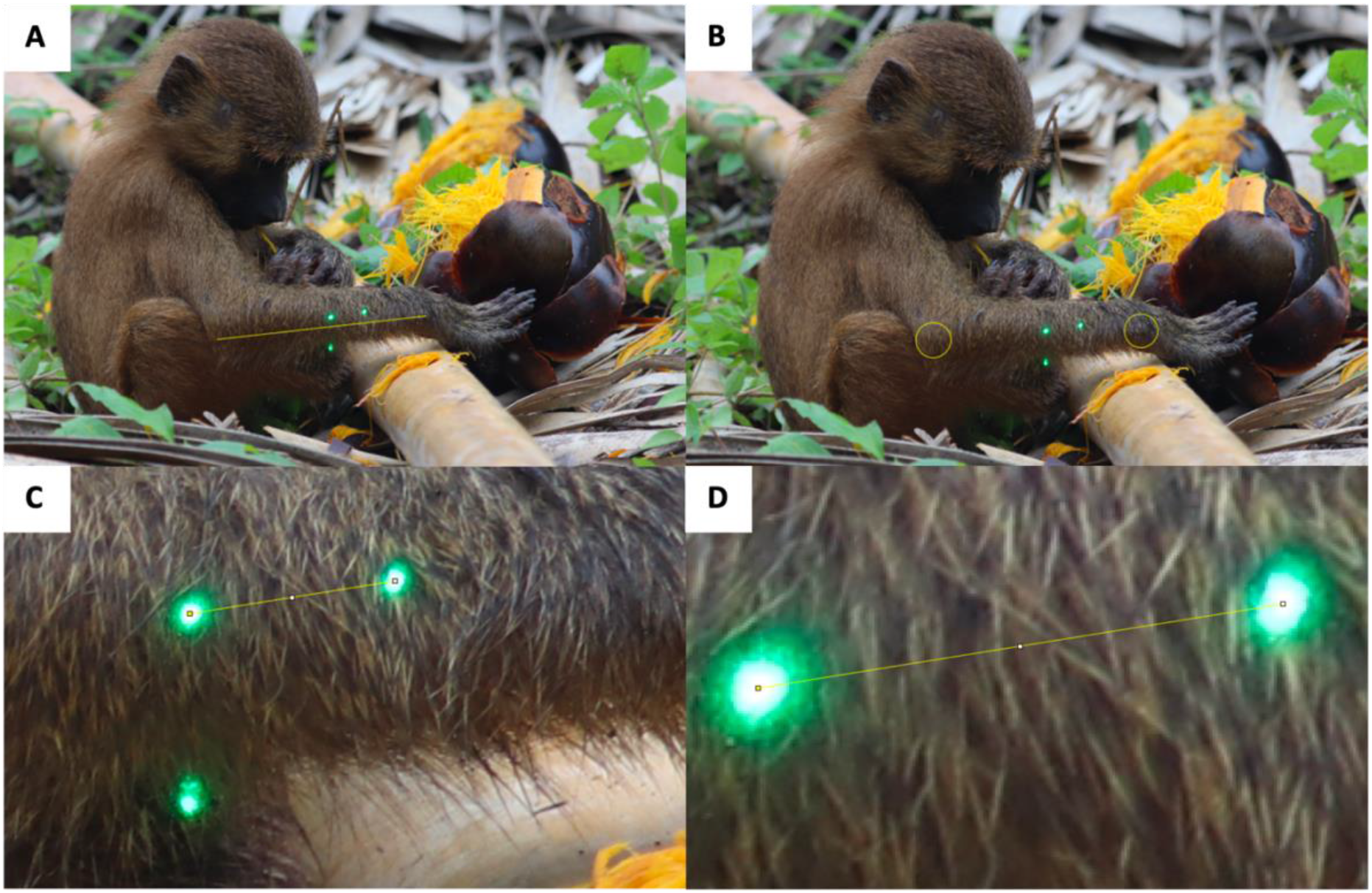
Lower arm measurement (A) in infant Guinea baboons is determined by body landmarks: olecranon (B, left yellow circle) and ulnar styloid process (B, right yellow circle). The horizontal lasers were used as a scale (inter-beam distance: 20 mm). Inter-laser distance was measured in ImageJ from centre to centre of the green laser dots (C, D). In our study, only horizontal lasers were used, due to a calibration issue with the vertical lasers

To minimize error, when we selected pictures to be measured, as those which met the following criteria (adapted from Galbany et al., 2017): landmarks were identifiable in the picture, and in focus; individuals had to be either sitting or standing, but always flexing their arm (Fig. S3.2, A-B); an ideal position was one in which the individual had the upper and lower arm forming a right angle. Laser points were on the same plane as the body landmarks of the infant. This plane was visually perpendicular to the optical axis of the camera objective lens and to the projection of the laser beams, so that the body of the infant was not rotated. Further, preferably the infant was situated in the centre of the picture.

To control for possible irregularities in the laser alignment and because we could not adjust and calibrate the lasers (i.e., inter-beam distance) during the field season, we projected them onto a millimetre paper. We took pictures at 2 and 5 m before and after each field day on which we took pictures. The pictures were later measured with ImageJ. This way, we could know the exact daily inter-laser distance and, therefore, the exact scale, which we used as a daily scale, since we had calibration problems during the study period: over time, the vertical lasers deviated from their original distance considerably, while the horizontal ones deviated from it only slightly. The vertical lasers were ultimately not used for measuring growth, as the differences were too large when increasing the distance between the photographed object and the camera. The calibration issues might have occurred because the apparatus was carried during field work and occasionally walked through dense terrain or due to temperature changes, since the inter-laser distance was shorter at the peak of the dry, hottest season for two years.

### S4. Mother-infant behaviours

**Table S4.** Fitted values of the GLMMs for the eight behaviours that showed variation in maternal style. These values represent the population average behaviours throughout the infancy period (from birth to 18-months of age). For maternal grooming, contact-sit, carry, time 1m proximity, and nipple contact fitted values indicate proportion of time; for maternal approaches, maternal leaves, maternal protections, and maternal rejections they indicate number events per hour

|  | Month |  |  |  |  |  |  |  |  |  |  |  |  |  |  |  |  |  |  |
| --- | --- | --- | --- | --- | --- | --- | --- | --- | --- | --- | --- | --- | --- | --- | --- | --- | --- | --- | --- |
| Behaviour | 0 | 1 | 2 | 3 | 4 | 5 | 6 | 7 | 8 | 9 | 10 | 11 | 12 | 13 | 14 | 15 | 16 | 17 | 18 |
| Maternal grooming | 0.022 | 0.022 | 0.022 | 0.023 | 0.023 | 0.023 | 0.023 | 0.023 | 0.023 | 0.023 | 0.024 | 0.024 | 0.024 | 0.024 | 0.024 | 0.024 | 0.024 | 0.025 | 0.025 |
| Contact-sit | 0.503 | 0.467 | 0.430 | 0.395 | 0.360 | 0.327 | 0.296 | 0.266 | 0.239 | 0.213 | 0.190 | 0.168 | 0.149 | 0.131 | 0.115 | 0.101 | 0.089 | 0.077 | 0.068 |
| Carry | 0.415 | 0.381 | 0.349 | 0.318 | 0.289 | 0.261 | 0.236 | 0.211 | 0.189 | 0.169 | 0.150 | 0.133 | 0.118 | 0.105 | 0.092 | 0.081 | 0.071 | 0.063 | 0.055 |
| Time 1m proximity | 0.875 | 0.849 | 0.819 | 0.784 | 0.745 | 0.702 | 0.654 | 0.603 | 0.550 | 0.496 | 0.442 | 0.389 | 0.338 | 0.291 | 0.249 | 0.210 | 0.176 | 0.147 | 0.121 |
| Nipple contact | 0.495 | 0.457 | 0.420 | 0.383 | 0.348 | 0.314 | 0.283 | 0.253 | 0.225 | 0.200 | 0.177 | 0.156 | 0.137 | 0.120 | 0.105 | 0.092 | 0.080 | 0.069 | 0.060 |
| Maternal approaches | 0.013 | 0.013 | 0.013 | 0.013 | 0.012 | 0.012 | 0.012 | 0.012 | 0.012 | 0.012 | 0.011 | 0.011 | 0.011 | 0.011 | 0.011 | 0.011 | 0.011 | 0.010 | 0.010 |
| Maternal leaves | 0.022 | 0.023 | 0.025 | 0.026 | 0.028 | 0.030 | 0.032 | 0.034 | 0.037 | 0.039 | 0.042 | 0.045 | 0.048 | 0.051 | 0.054 | 0.058 | 0.062 | 0.066 | 0.071 |
| Maternal protections | 0.047 | 0.030 | 0.019 | 0.012 | 0.008 | 0.005 | 0.003 | 0.002 | 0.001 | 0.001 | 0.001 | 0 | 0 | 0 | 0 | 0 | 0 | 0 | 0 |
| Maternal rejections | 0.002 | 0.002 | 0.002 | 0.002 | 0.002 | 0.003 | 0.003 | 0.003 | 0.003 | 0.004 | 0.004 | 0.004 | 0.005 | 0.005 | 0.006 | 0.007 | 0.007 | 0.008 | 0.009 |

### S5. Factor analysis

**Figure S5.**
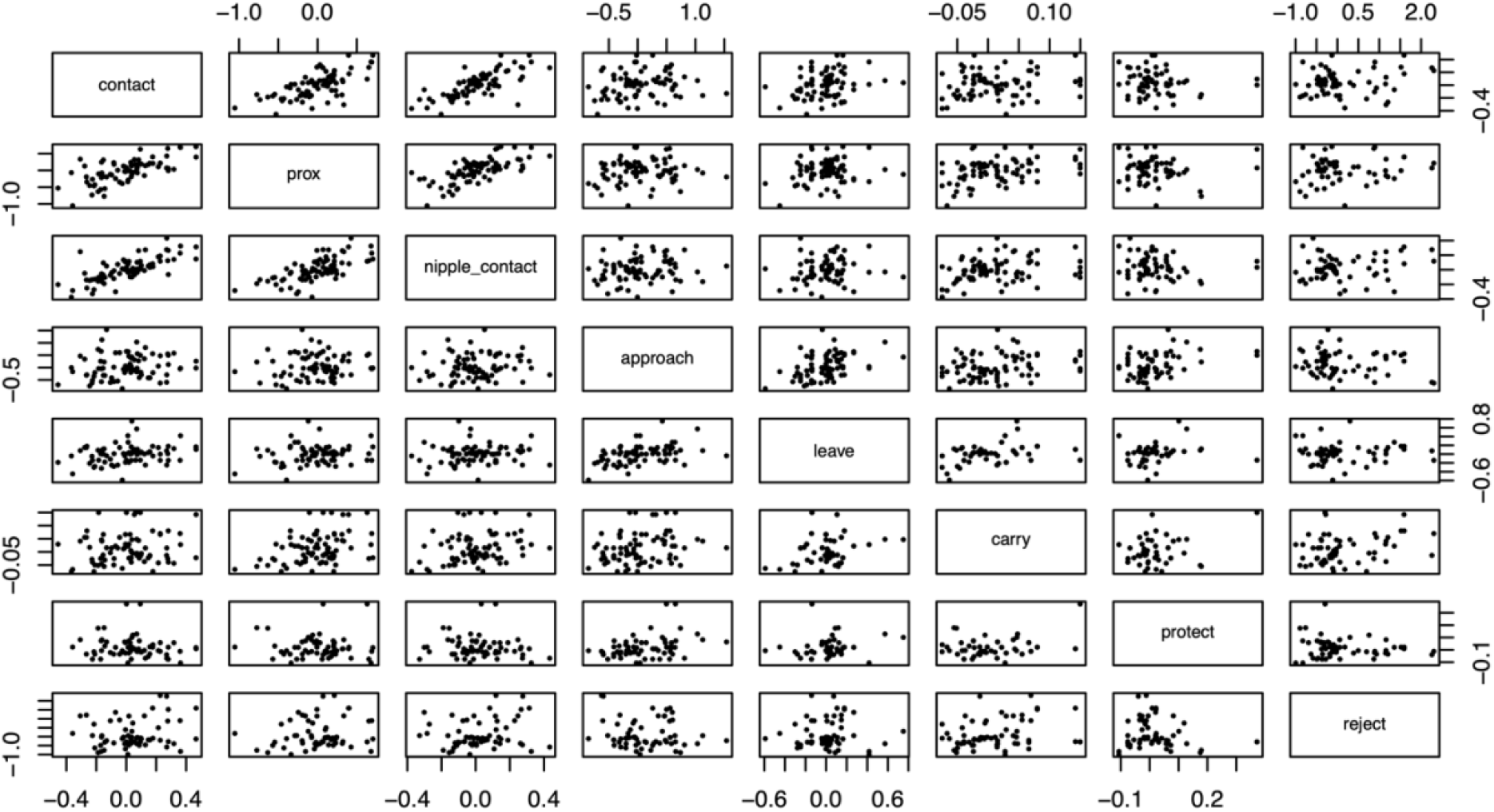
Scatter plot depicting the correlations between the eight behaviours that showed sufficient variation between mothers (i.e., >0.1 Standard Deviation). Each point represents the estimated BLUP for a given mother-infant pair

**Table S5.**
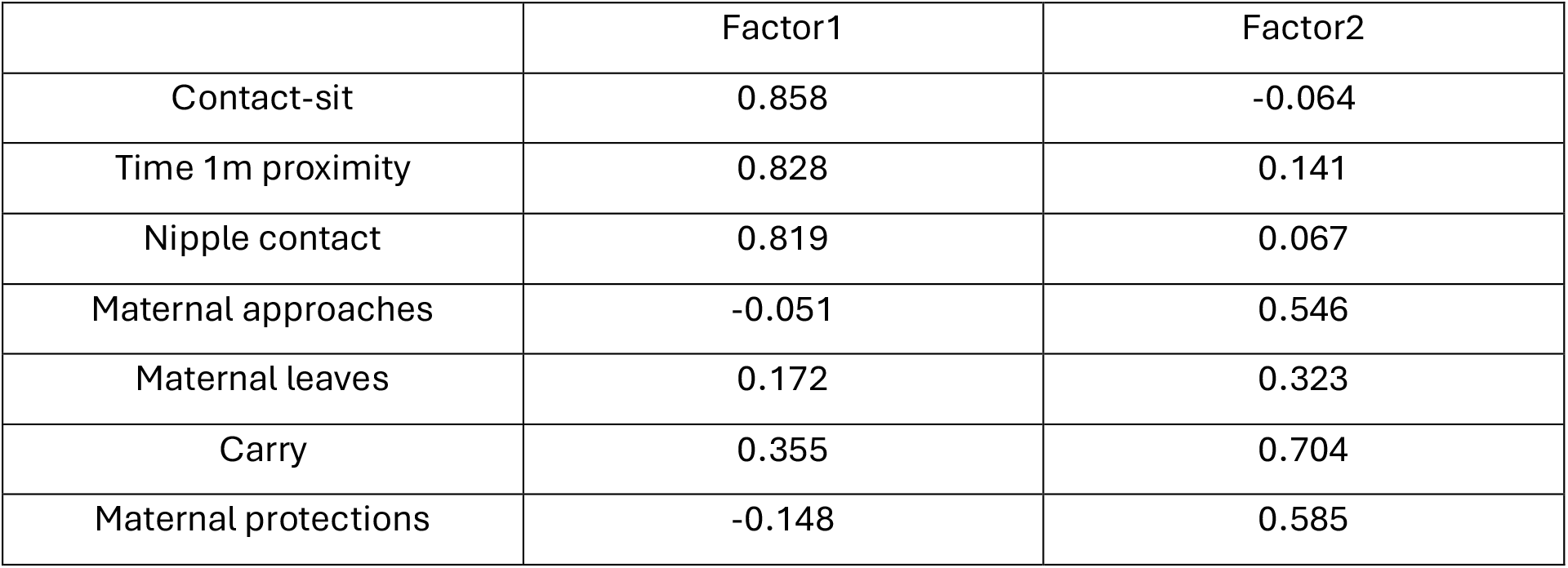
Table with detailed factor loadings of the considered behaviours on the two-factors

|  | Factor1 | Factor2 |
| --- | --- | --- |
| Contact-sit | 0.858 | -0.064 |
| Time 1m proximity | 0.828 | 0.141 |
| Nipple contact | 0.819 | 0.067 |
| Maternal approaches | -0.051 | 0.546 |
| Maternal leaves | 0.172 | 0.323 |
| Carry | 0.355 | 0.704 |
| Maternal protections | -0.148 | 0.585 |

### S6. Ecological conditions (NDVI)

We controlled for environmental conditions due to the potential effect of food availability on the infant survival and growth. The Normalized Difference Vegetation Index (NDVI) was used as a proxy for food availability. The NDVI is a widely used bio-geophysical indicator of the land’s green plant coverage - or vegetation density - and, therefore, it is considered an indirect measure of rainfall. The NDVI values were obtained from the Copernicus Global Land Service (CGLS) (https://land.copernicus.eu/global/), which is part of the Earth Observation programme of the European Commission. We downloaded the NDVI values at 300 m spatial resolution from 2016-2022 (accessed in July 2022); although the behavioural data for the infants of the study were first collected in 2017, we included NDVI data since 2016 because some of the infants in the survival model were born during that year. Values of NDVI were based on a 10-day maximum composite value derived from daily acquisitions of Earth land surface reflectance observations from the sensor PROVA-B (product version 1, from 2016-2020: Swinnen & Carolien, 2016) and sensor Sentinel-3 OLCI (version 2, from 2021: Swinnen & Carolien, 2022).

We limited values of NDVI to our study region (Lat 13°0’14.252’’ to 13°5’39.647’’; Long - 13°13’24.813’’ to -13°18’54.438’’) and used the function ncvar_get of the package ncdf4 (version 1.19: Pierce, 2021) in R (R Core Team 2022) to extract the NDVI values from the downloaded files. We excluded non-optimal observations (digital value >250), which created interferences due to their origin in cloud coverage and ocean. In case of missing NDVI values, we computed a linear interpolation between the last and next days for which we had available data (Fig. S6). The daily NDVI values were averaged across the study area and later used as a control fixed effect for our main growth and survival models.

**Figure S6.**
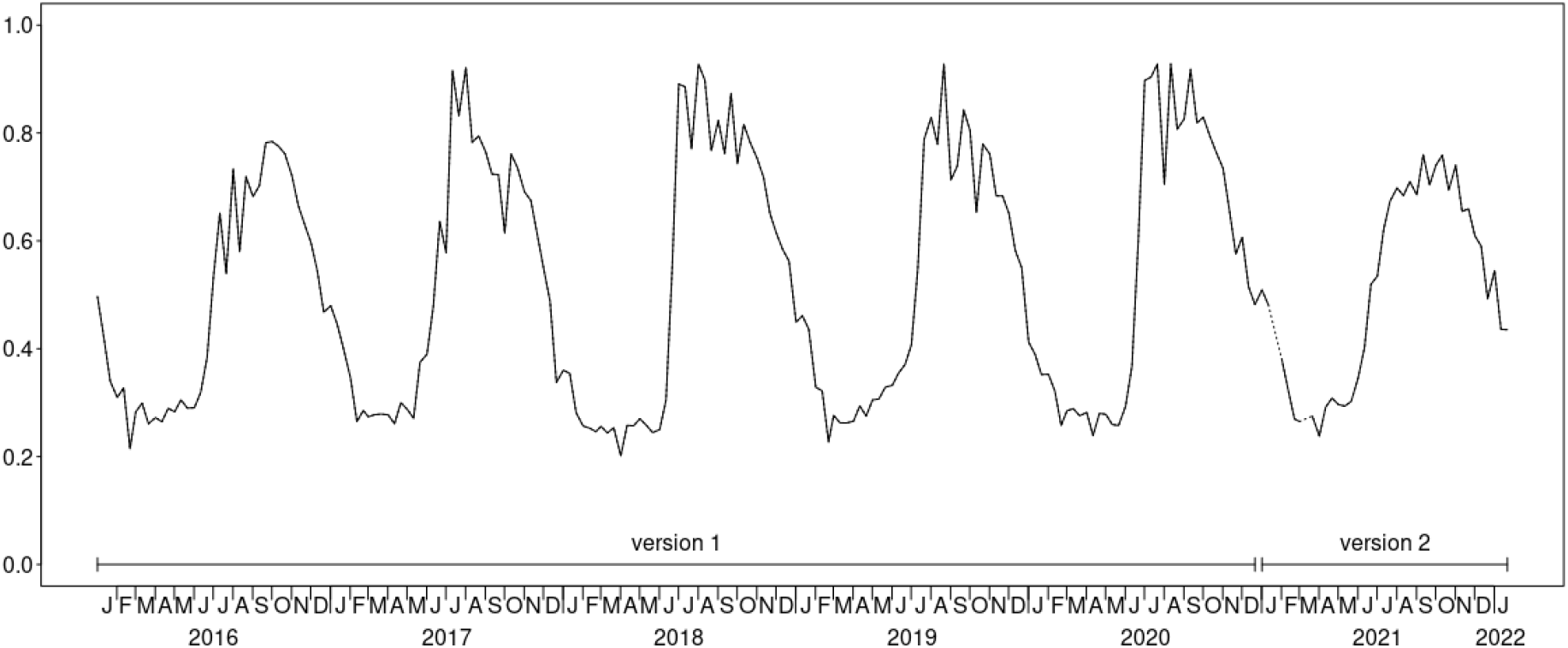
NDVI values in the study region during the study period extracted from versions 1 and 2 of the Copernicus Global Land Service (CGLS). Note the pronounced seasonal variation in plant productivity, being lowest in the driest months (March-May) and most pronounced in the rainy season (June-October). The dashed line represents interpolated values.

### S7. Maternal style consistency for mothers with multiple infants

**Figure S7.1.**
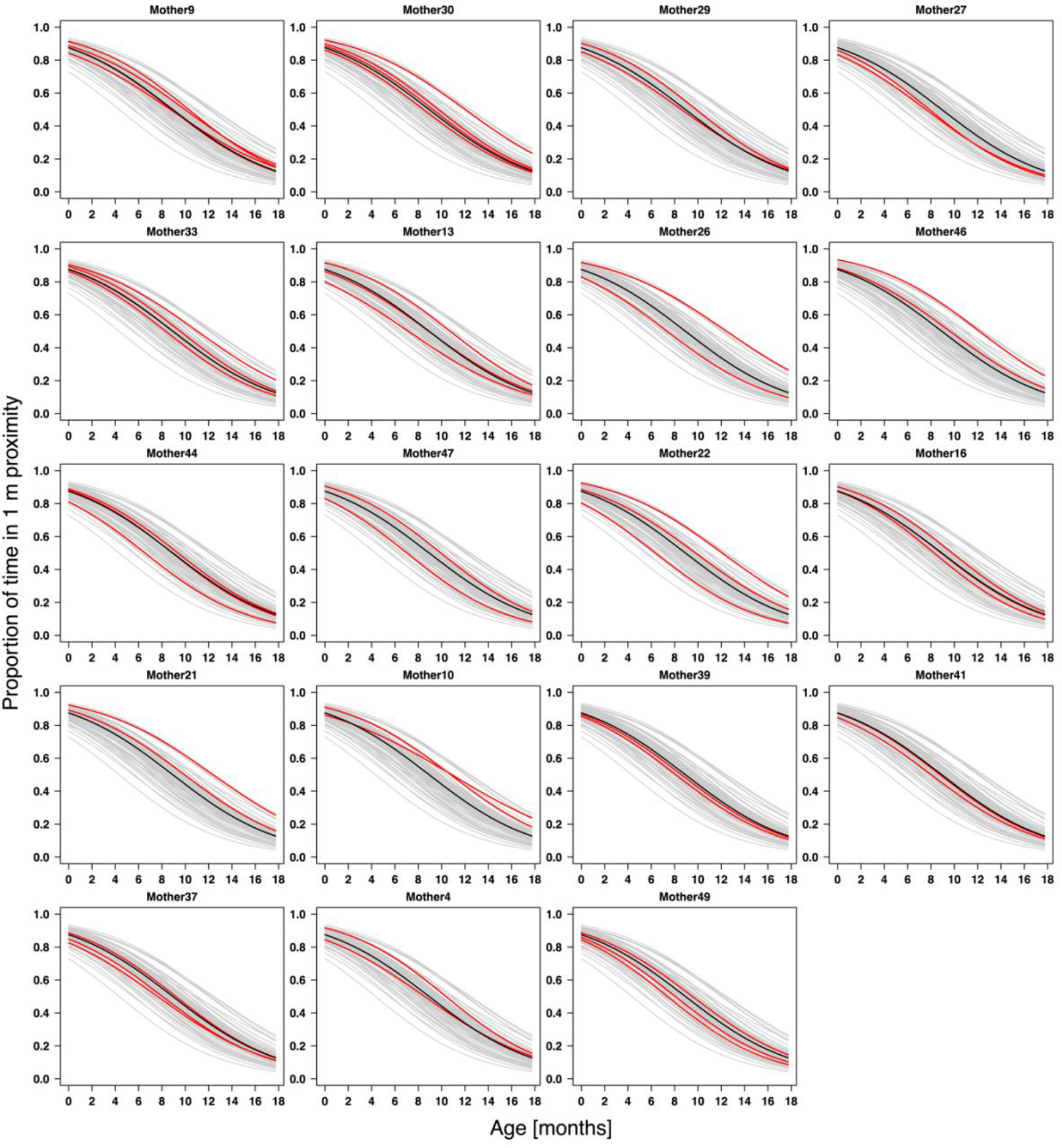
Predicted proportion of time in 1-m proximity across age for mothers with multiple infants (N = 19). Red lines show predicted trajectories for each mother and her infants, grey lines represent all other mother-infant combinations, and the black line represent the population mean.

**Figure S7.2.**
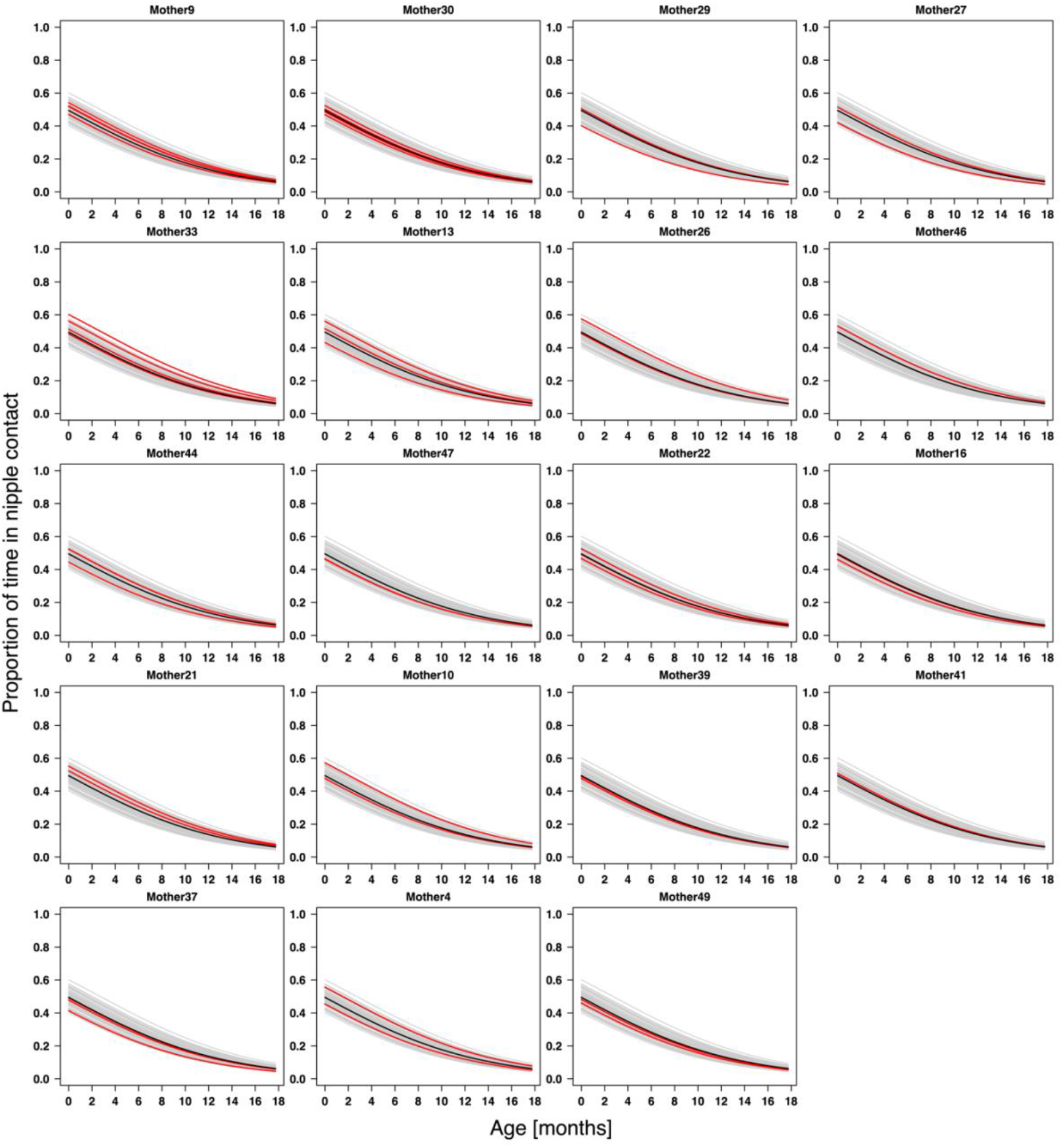
Predicted proportion of time in nipple contact across age for mothers with multiple infants (N = 19). Red lines show predicted trajectories for each mother and her infants, grey lines represent all other mother-infant combinations, and the black line represent the population mean.

**Figure S7.3.**
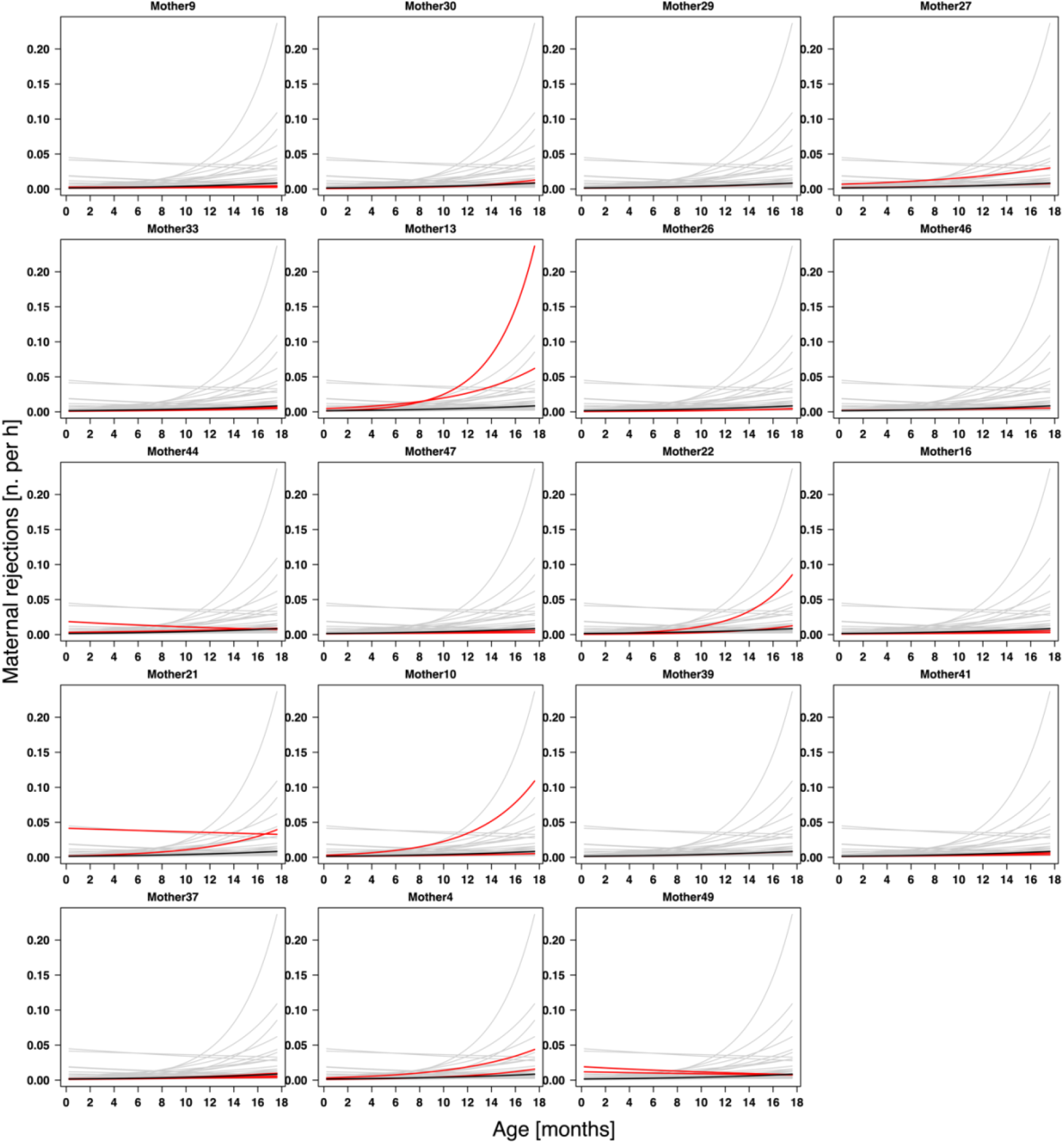
Predicted number of maternal rejections per hour across age for mothers with multiple infants (N = 19). Red lines show predicted trajectories for each mother and her infants, grey lines represent all other mother-infant combinations, and the black line represent the population mean.

### S8. Repeatability of maternal behaviour

In this section we report the complete methods used to assess the repeatability of maternal behaviors, while in the main draft we only provide a summary.

Following a reviewer’s suggestion, we formally assessed the repeatability of maternal behaviour using adjusted intraclass correlation coefficients (ICC; Nakagawa et al., 2017). A complication arose because our models included random slopes of infant age within mother ID (and infant ID). In such models, the estimated variance associated with the random intercept of mother ID is conditional on infant age being zero. Since infant age was mean-centred in all models, the estimated variance of the mother intercept corresponds to the average infant age in the dataset. However, the average age in our dataset was influenced by irregularities in longitudinal sampling, including temporary or permanent disappearance of infants, interruptions caused by the COVID-19 evacuation, and other logistical constraints. Consequently, directly calculating ICCs from the fitted models would not yield biologically meaningful estimates of repeatability.

We therefore applied two different approaches depending on the results of the fitted model. For behaviours in which the random slope of infant age within mother ID was estimated to be negligible (< 0.001), we calculated ICCs directly from the fitted models. This approach was applied to contact-sit, carry, nipple contact, protection, and proximity. For behaviours in which the random slope of infant age within mother ID was non-negligible (approach, reject), we first generated fitted trajectories across the observed infant age range using the fixed effects estimates together with mother- and infant-specific deviations (i.e., Best Linear Unbiased Predictors; BLUPs; Baayen, 2008). We then quantified, for each mother–infant dyad, the average deviation between these dyad-specific fitted values and the fitted values expected for an average mother with an average infant across the infant age range in our data (based solely on the fixed effects component; see also Fig. SI 25 in Mundry et al., 2023). These deviation estimates were subsequently analysed using linear mixed-effects models containing only an intercept as fixed effect and mother ID as a random intercept. ICCs derived from these models were used as estimates of repeatability. This procedure was applied to maternal approaches and rejections.

Neither approach could be applied to maternal leaves. In this model, the random slope of infant age within mother ID was relatively large (estimated SD = 0.44), whereas the infant-level random effects were essentially zero. As a consequence, estimated trajectories were effectively identical across infants within mothers, resulting in a residual variance approaching zero and causing convergence problems in the subsequent linear mixed-effects model.

### S6. Effect of maternal age and infant sex on maternal style

**Table S9.**
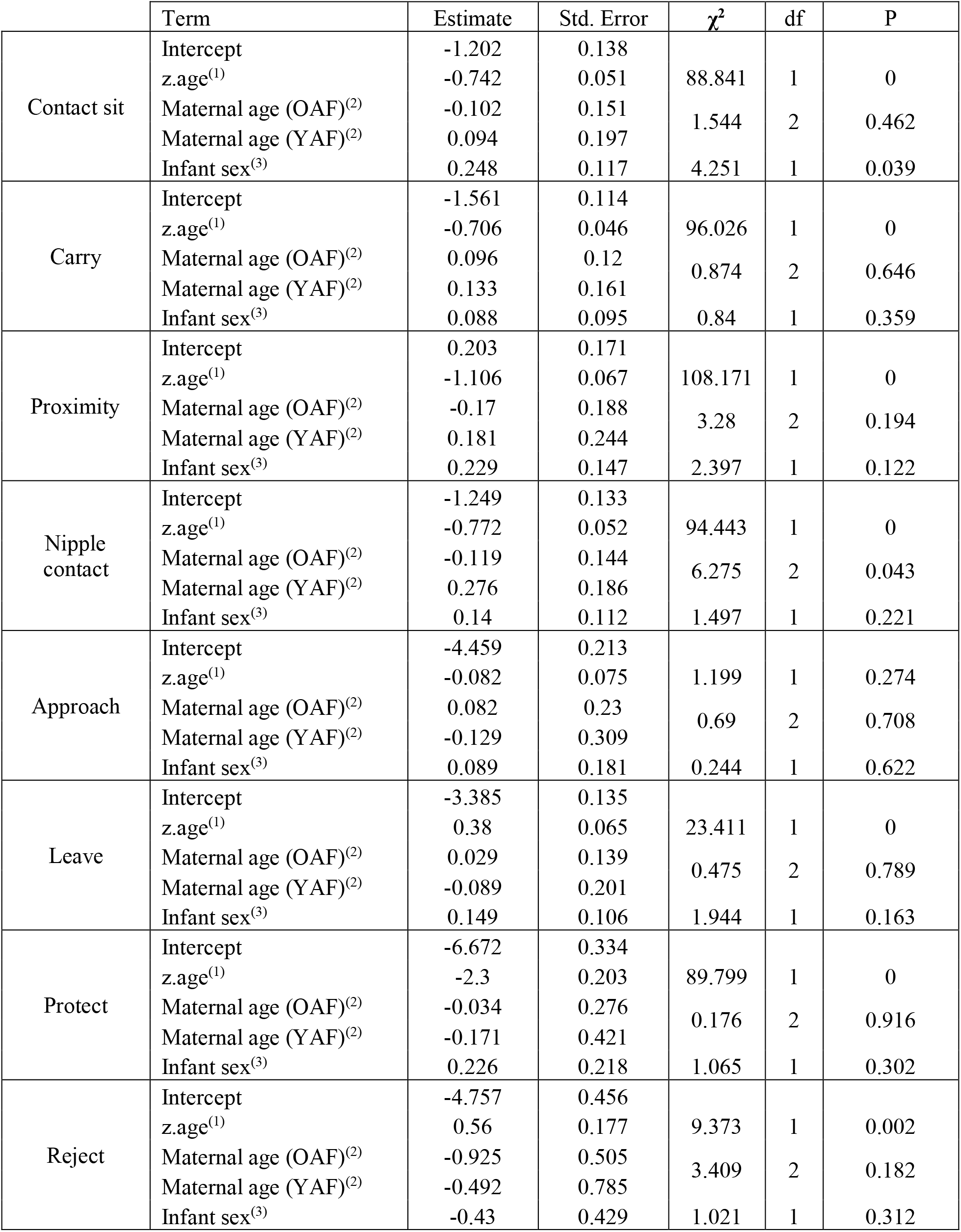
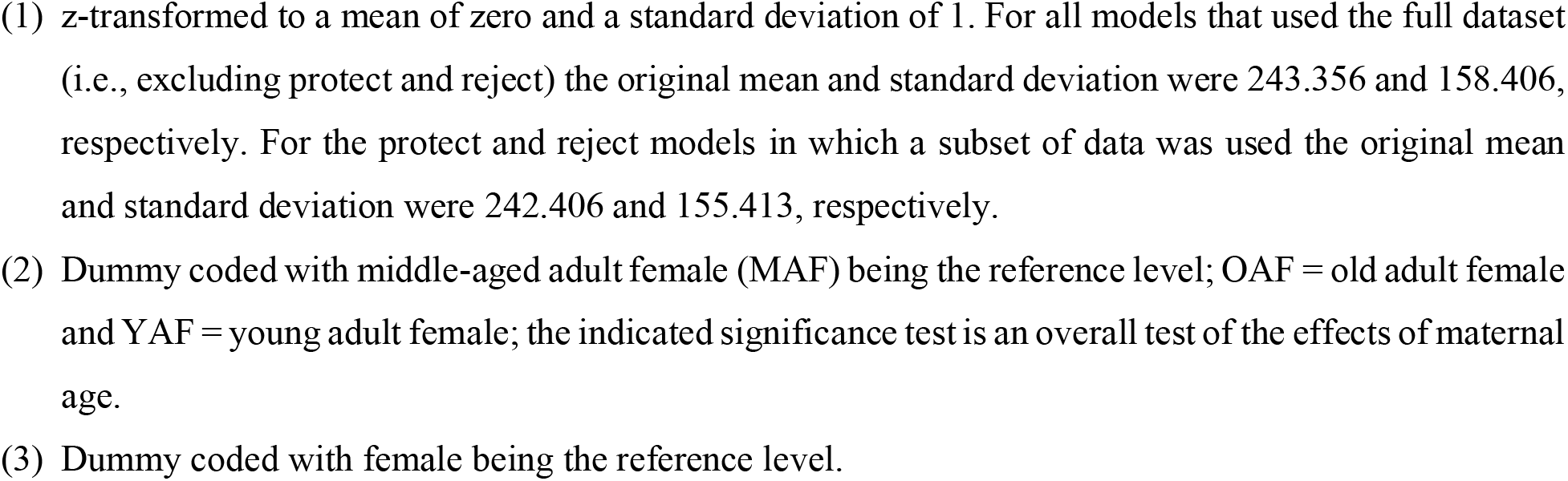
Table with detailed results of the models testing the effect of maternal age and infant sex on maternal style (one model per maternal behaviour)

